# A unified model of hippocampal spatial and object cells involving bidirectionally coupled Lateral and Medial Entorhinal Cortical layers

**DOI:** 10.1101/2024.09.09.612040

**Authors:** Bharat K. Patil, Azra Aziz, Manoj Kumar Saka, Sachin S. Deshmukh, V. Srinivasa Chakravarthy

**Affiliations:** Computational Neuroscience Lab, Indian Institute of Technology Madras, Chennai, India; School of Natural Sciences, Shiv Nadar University, Delhi NCR, India; Institute of Science and Technology Austria, Vienna, Austria

## Abstract

Popularly referred to as the GPS of the brain, the hippocampus has a variety of neurons that encode spatial properties of the environment. These spatial cells of the hippocampus may be broadly placed under two categories – those that encode spatial locations (e.g. place cells, grid cells etc) and those that encode spatial objects (eg. Object-sensitive cells. Object-trace cells etc). There are computational models that explain emergence of specific types of spatial cells, but it is challenging to construct integrative models that can demonstrate the emergence of the complete range of spatial cells both space and object type. We present a simple, unified computational model that explains the emergence of a wide variety of object- and spatially-sensitive neurons in the hippocampus. The model is essentially a deep neural network that combines visual and path integration information. The visual information is received by a part of the model that is analogous to Lateral Entorhinal Cortex (LEC) and path integration information is received by a layer analogous to Medial Entorhinal Cortex (MEC). In order to arrive at a consistent estimate of position, LEC and MEC in the model are connected laterally using a Graph Neural Network. The model is trained to predict position, orientation and reward of a simulated agent. The agent explores a box-like environment with colored walls and objects on the floor and is rewarded based on its encounters with objects. The model demonstrates the emergence of the following 7 types of spatial and object cells - place, grid, border, object, object-sensitive, object-vector and, object-trace cells. The model findings compare favorably with a large body of experimental literature on hippocampal spatial cells.

## Introduction

Real-world environments present a great diversity of objects, which often serve as valuable landmarks aiding navigation and foraging in the environment (Gothard *et al*., 1996; Chan *et al*., 2012). Animals have demonstrated their ability to utilise these objects to navigate through different settings and reach specific locations. Popularly dubbed as the GPS of the brain, the hippocampus has been subjected to decades of investigation aimed at understanding the neural substrates of spatial environments and their contents. Researchers have delved into hippocampal formation to unravel the representations of objects in various environments. These investigations have provided valuable insights into the neural circuitry underlying object processing, leading to the discovery of diverse cells including object cells that fire in the vicinity of the objects (Tsao, Moser and Moser, 2013), object vector cells whose firing field maintains a vector relationship with the object at multiple locations, for various shapes and sizes of the object (Høydal *et al*., 2019), and object-sensitive cells whose firing field has more spatial information in presence of objects (Deshmukh and Knierim, 2011) in the Entorhinal Cortex, along with landmark vector cells, in the hippocampus proper, that fire at a particular distance and direction from object or objects (Deshmukh and Knierim, 2013). These findings illuminate the essential role of neural substrates in the spatial mapping of objects within the hippocampal formation.

Additionally, an extensive body of literature (Young *et al*., 1997; Clark *et al*., 2002; Gilbert and Brushfield, 2009; Girardeau *et al*., 2009) has presented compelling evidence supporting the hippocampus’s involvement in memory processing. A subset of experiments (Deshmukh and Knierim, 2013; Tsao, Moser and Moser, 2013) focused explicitly on object memory within the hippocampal formation, exploring both spatial and non-spatial contexts. Such research provides valuable insights into the intricate mechanisms of object memory and its interplay with spatial representations within the Hippocampus.

A plethora of modelling approaches have been taken to model the subset of these spatial neurons. Hetherington and Shapiro (Hetherington & Shapiro, 1993) proposed an Elman recurrent neural network with hidden layer that produced place-cell-like activity. O’Keefe and Burgess (O’ Keefe and Burgess, 1996) proposed a model that describes the properties of place fields based on environmental geometry. Li et al. (Li et al., 2020) proposed a multisensory integration neural network to model motion-based place cells (MPCs), vision-based place cells (VPCs), and conjunctive place cells (CPCs). Solstad et al. (Solstad et al., 2006) describe a model that can produce place cells from grid cells. There are two approaches to modelling grid cells: the oscillatory interference models (Burgess, Barry and O’Keefe, 2007; Giocomo and Hasselmo, 2008; Zilli and Hasselmo, 2010) and, the continuous-attractor neural network models (Sargolini *et al*., 2006; Burak and Fiete, 2009). There are also some models that model subsets of place, grid and, border cells together (Cueva and Wei, 2018; Soman, Muralidharan and Chakravarthy, 2018; Aziz *et al*., 2022). While these models effectively address subsets of spatial cells, there is a pressing need for a simplified, comprehensive model that attempts to integrate all spatial and object-responsive cells into a single framework.

In addition to previous studies, the Tolman-Eichenbaum Machine (TEM) presents a bimodular model that uses medial entorhinal cells to form a structural knowledge base and hippocampal cells to link this base with sensory representations, thereby integrating spatial and relational memory. TEM effectively demonstrates diverse hippocampal functions, including spatial and landmark representations after learning. However, while TEM’s predictions regarding structural knowledge preservation align well with certain empirical observations, the model’s dependence on predefined graph structures simplifies the adaptive nature of the hippocampal-entorhinal system, confining it to noise-free environments. Furthermore, the model’s input facilitates exclusion of sensory correlates in the environment to isolate transition structure, which diverges from real-world settings.

In this paper, we describe a unified model that describes a wide variety of object and spatially selective cells in the hippocampus. The model incorporates multisensory integration in a manner that is consistent with the anatomical correlates in Entorhinal Cortex (EC). The model is essentially a deep neural network that combines two sensory pipelines using a special form of lateral connectivity in one of the hidden layers. The two sensory pipelines that are combined are 1) the path-integration (PI) pipeline from locomotion inputs and 2) the visual processing pipeline from visual inputs. Information from these two pipelines is combined using a Graph Convolution Network (GCN) that represents the lateral connections between the Lateral Entorhinal Cortex (LEC) and Medial Entorhinal Cortex (MEC) (Kipf and Welling, 2016). Beyond the GCN level, the network contains three fully connected layers that are reminiscent of the CA regions of the Hippocampus. By combining PI and visual inputs, the network is trained to estimate the following variables: the current position (x, y), the current heading direction (Θ), and the expected reward (r), set as a reward of +1 when the agent is near the object and 0 elsewhere. As we will show below, a wide variety of object cells emerge naturally in the network trained in the aforementioned fashion.

The paper is structured as follows. The Methods section begins with the generation of input data, which includes trajectories and corresponding visual data. The data is generated using from Unity software. It then provides a detailed explanation of the model architecture and the training procedures. The Results section first introduces a unified model for spatial and object-responsive cells, followed by an in-depth analysis of each cell type under various environmental and training conditions. The Discussion section summarizes the key observations and findings from the study, offering insights and implications for future research. The proposed comprehensive and simple model thus developed shows emergence of a variety of spatial cells (place, grid, border, object, object-sensitive, object-vector and, object-trace cells) observed in the Hippocampal formation.

## Methods

The current study aims to model a multitude of object representations (object-sensitive, object cells) and neuronal vector coding (object-vector cells) along with place, grid, border and object-trace cells from numerous experiments using a single comprehensive model. To this end, using Unity3D, we designed a virtual environment inspired by actual experiments. A point-sized simulated agent navigates inside the environment on a random 2D trajectory. From every point on the trajectory, the agent gets a view of the environment, as seen in the forward direction along the tangent to the trajectory. Thus, every location and direction on the trajectory is associated with an image corresponding to the view. The self-motion information and the stream of views thus generated are used to train the proposed model.

### Trajectory generation for 2D environment

The agent is allowed to navigate through a square environment of edge size 2 units. This is executed by generating a random sequence of points, while imposing constraints on collisions, sudden turns, and speed (Soman, Muralidharan and Chakravarthy, 2018) (Supplementary section: 13). This sequence of points is then interpolated and smoothened using a “spline” algorithm (Fig. 1a). When an object is introduced into the environment, we constrain the trajectory so that no part of the trajectory passes through the space occupied by the object. We use the smoothened trajectory points to generate movement information, such as the heading direction (same as the movement direction for this model) and position. To model the experiments mentioned above, specific modalities from individual experiments, such as an introduction or shifting of objects, boundaries, and cues, have been incorporated to simulate the trajectories.

**Fig 1:**
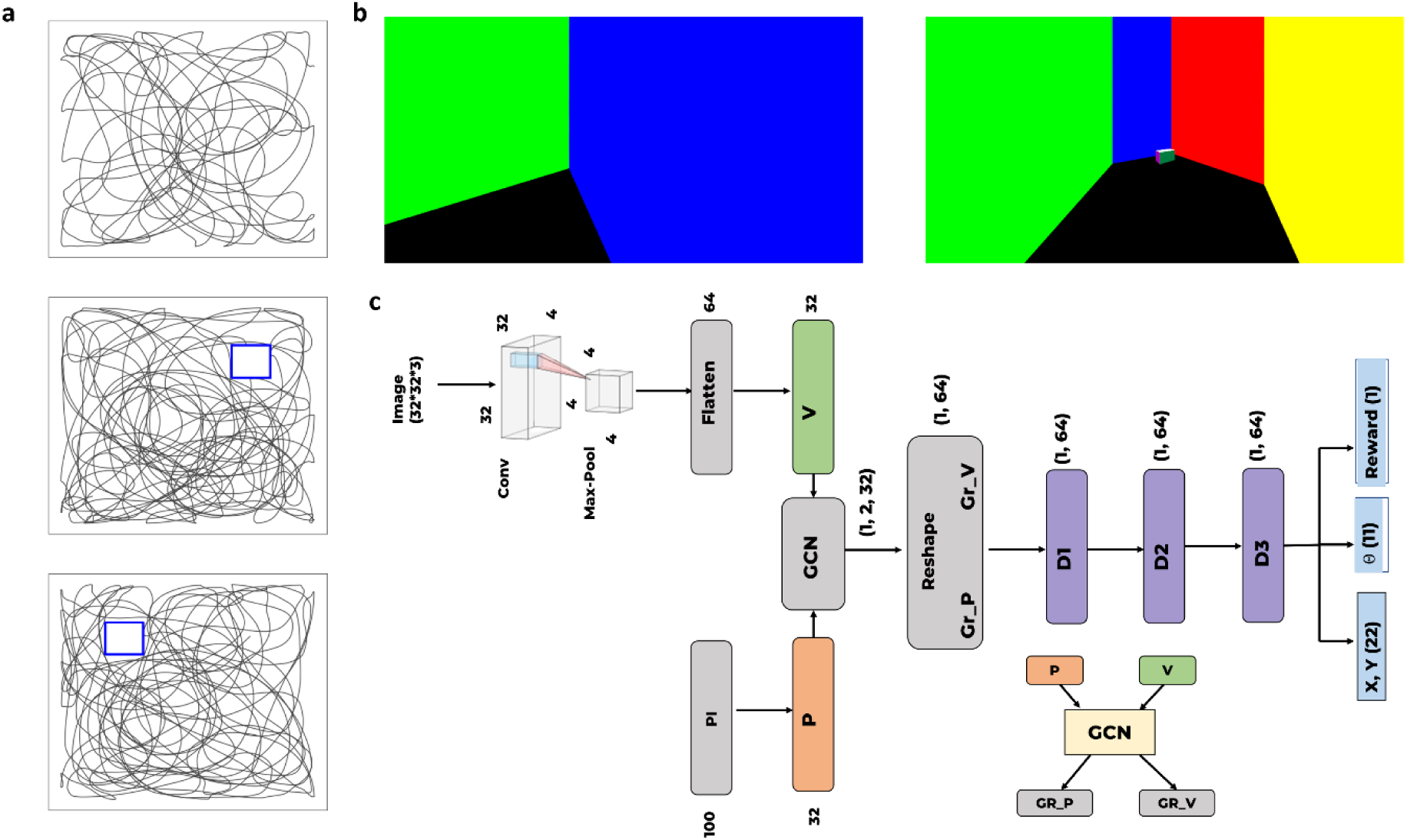
a) Trajectories, Top: session 1 (without object). Centre: session 2 (with object). Bottom: session 3 (with shifted object). b) snapshots of the environment from Unity software showing examples of views seen by the agent. Right shows an object in the environment. c) Model architecture showing visual and PI pipelines. These pipelines are convolved using the Graph Convolution Network (GCN) layer (subpanel), post which three fully connected layers are present, and the model predicts position (x, y), direction (θ), and reward.

### Virtual Reality Environment

We designed a virtual box environment for visual input using Unity3D, keeping the dimensions consistent with the trajectory bounds (edge length for the surface/floor is 2 units). The agent’s point of vision (POV) (originating at the eye level in animals) is at a height, ‘h’ from the surface (Fig. 1b). The height of the walls (H) is substantially larger than the height of the agent’s POV, so that no distal cues other than the colors on walls are visible. The field of vision for the agent is set at 120°. For most of the simulations, the walls of the rectangular environments are distinct in colour that serve as distal cues and the surface is dark (black) unless specified otherwise.

A subset of experiments involves introducing novel objects, displacing objects to a novel location, or removing the object from the environment. For these experiments, a cubical object is used with edge length 0.3 units and distinct colours on all faces unless specified otherwise. The agent’s POV image (RGB), generated at every point on every trajectory, has a size of 1171*430*3 and further rescaled to 32*32*3, which serves as input to the visual pipeline of the model.

### Firing Field Detection

The activation for each neuron is recorded for an entire trajectory for each session in all experiments. A threshold value of 1.5 standard deviations higher than the mean firing rate is applied to each individual neuron. These thresholded values are then used to generate firing rate maps that are visually inspected to classify the cells further.

## Model

The proposed model consists of a PI pipeline and a Visual Pipeline which are combined so as to achieve a unified spatial representation of the environment (Fig. 1).

### Path Integration Pipeline

Path Integration (PI) is implemented as an array of sinusoids whose frequency is modulated by velocity as described in (Aziz *et al*., 2022) (Supplementary section: 9). This involves a Head Direction (HD) layer connected one-to-one with the PI array (Fig. 1). The HD layer has 100 neurons, each with its unique preferred direction. The HD layer is modelled as:

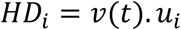

where, *v*(*t*) is the velocity of the animal in the 2D plane and *u*_*i*_, a 2D unit vector, is the preferred direction of the i^th^ HD neuron.

The PI layer is modelled as an oscillatory array of neurons, the output of which is further time-averaged to remove time dependency and make the responses of PI neurons purely functions of space (Aziz *et al*., 2022) (supplementary section: 9). The response of the PI layer is modelled as,

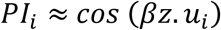

where z is the displacement vector of the agent, and *β* is the modulation factor.

The output of the PI layer is then passed through a fully connected layer so that the visual and the PI pipeline are equidimensional.

### Visual Pipeline

Visual Pipeline for the model is designed as a Convolutional Neural Network (CNN) that consists of a pair of convolution and max pooling layers. The input to the convolution layer is a stream of POV images of the agent with RGB channels, which are captured when the agent follows the stipulated trajectory. The output of the final max pooling layer is then flattened and passed through a fully connected layer (Fig. 1c).

### Graph Convolution Network

A Graph Convolution Network (GCN) (Kipf and Welling, 2016) models the lateral connections between the outputs of both the pipelines (Fig. 1c). The layer activations from both pipelines behave as nodes for the Graph Neural Network (GNN) (Scarselli *et al*., 2009), where each element of a specific layer is assumed to be a feature of that node. Hence, the layer’s dimension equals the feature length for all nodes of the GNN. The current model’s GNN layer receives inputs from two layers (PI and Vision), with 32 features for each. The output of each node of the GNN is given as:

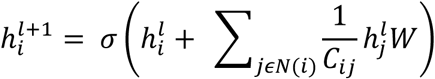

where, *h_i_*^*l*^: feature vector of i^th^ node before convolution; *h_i_*^*l*+1^: feature vector of i^th^ node after convolution; W: weight matrix; *C*_*ij*_: degree of node, N: number of nodes.

The GNN weights act as lateral connections observed between LEC and MEC in Hippocampal formation that holds primacy for the emergence of object-based and vector-based representations.

The output of GCN is flattened and is further passed through 3 fully connected layers, each of dimension 64, that were used to predict instantaneous reward, current position, and heading direction.

For efficient learning and increased accuracy, the agent’s position (x and y coordinates) and heading direction are predicted as a population code. The agent’s spatial coordinates are represented as a Gaussian with its mean as the correct coordinate value and a 0.3 standard deviation around it to generate the population code for the position. Hence the predicted current position is given as a set of two Gaussians, one centred at the correct x and another centred at the correct y coordinate of the agent. To generate population code for heading direction, we define a unit vector (ui) of length 11 in x and y components spanning 360°.

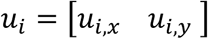

The instantaneous direction is given by:

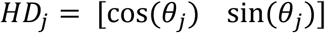

Therefore, the population code for Head Direction is defined as:

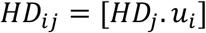

The model’s output is a 34-dimension array where the first 22 nodes predict the agent’s position (11 each for both coordinates), the next 11 nodes predict the heading direction, and the last node predicts the instantaneous reward.

### Training Procedure

Training procedure and environment vary depending on the neuron group being modelled. The model is first prepared by training on an environment with different coloured walls and without objects. The model is further trained by introducing objects into the environment under various conditions. Alterations are performed by adding or shifting an object, changing environment and object size, and changing environment or object attributes. Then depending on the experiment, the model is either trained or tested. The model representations exhibit a specific neuron family depending on the configuration employed separately for the position, heading direction, and reward (Table 1). Individual procedures are explained further under each cell type’s heading. From now on the training configurations for our models are represented using a binary encoding scheme, denoted as “config”, to specify the presence or absence of training on particular outputs. Each configuration is encoded as a three-digit binary string, where each digit represents whether the model is trained on a specific output: the first digit represents position, the second represents head direction, and the third is for reward. For instance, a ‘config 001’ indicates that the model is trained exclusively on reward, with the output layer only incorporating the reward component. Similarly, a config value of 101 signifies that the model is trained on both position and reward, but not on head direction (Table 1). Varying the configurations is shown to have a significant impact on the neural representations generated in the network. All training is performed by supervised learning.

**Table 1:** Configuration description for the training of the model.

| Configuration |  |  |  |
| --- | --- | --- | --- |
| Position | Head direction | Reward | Description |
| 0 | 0 | 1 | Model trained only for reward. |
| 0 | 1 | 0 | Model trained only for Head direction. |
| 1 | 0 | 0 | Model trained only for Position |
| 1 | 0 | 1 | Model trained only for Position and Reward. |
| 1 | 1 | 0 | Model trained only for Position and Head direction. |
| 0 | 1 | 1 | Model trained only for Head direction and Reward. |
| 1 | 1 | 1 | Model trained for Position, Head direction and Reward. |

## Results

We train and study the agent’s behavior in various virtual reality environments inspired by experimental paradigms. The model is trained to predict either position, heading direction, reward, or a stipulated combination of the three. Post-training, the neuronal responses from different hidden layers are observed to reveal a correspondence to spatial and object cell responses from the literature (O’Keefe and Dostrovsky, 1971; Fyhn *et al*., 2004; Hafting *et al*., 2005; Deshmukh and Knierim, 2011, 2013; Tsao, Moser and Moser, 2013; Bjerknes, Moser and Moser, 2014).

The current modelling study demonstrates emergence of 7 types of neurons used to represent the environment during navigation. They are place, grid, border, object-sensitive, object, object-vector and object-trace cells.

First, we train a general model as discussed in model architecture with ‘config 111’ configuration of the output layer and analyse all the neurons from the hidden layers. Generally, we follow a common sequence of experiments to analyse the model responses. In this study, we run three training sessions. The first training session is done on a square environment with different colours on each wall and no objects (session 1) (Fig 1a, top). The trained model in the first simulation is then retrained after introducing an object in the same square environment (session 2) (Fig. 1a, centre). Then the model from session 2 is further retrained after shifting the object to a new location (session 3) (Fig. 1a, bottom).

After training session 1, we analysed 5 hidden layers in the model, i.e., D1, D2, and D3, each with 64 neurons, a hidden layer in the vision pipeline (V), a hidden layer in the PI pipeline (P) each with 32 neurons and, 2 GCN layer (Gr_V and Gr_P) each with 32 neurons (Fig. 1c). The neuron output data was converted into firing rate maps (supplementary section: 3) which were further analysed to classify the neuron’s firing into one of the following cell types, i.e., place, grid, border, boundary vector, object sensitive, object, object vector, and object-trace cells.

To classify a neuron firing as place cell firing, spatial information (Skaggs *et al*., 1994) (supplementary section: 5) and sparsity score (supplementary section: 4) is calculated. The neurons whose spatial information > 0.3 bits/spike and have a sparsity score of < 0.1 are selected. Since these metrics are sensitive to low firing rates, we conducted a random shuffling test (supplementary section: 10) for these filtered cells. Furthermore, cells with spatial information and sparsity scores higher than the 95^th^ percentile of their individual distributions were labelled as place cells. Neurons with a Hexagonal Grid Score (HGS) greater than 0.1 were filtered as potential grid cells. Another random shuffling test (supplementary section:10) was performed to identify cells with an HGS greater than the 95^th^ percentile of the resulting distribution. These cells were labelled as grid cells. To identify border cells, a border score was defined similarly to the method described in (Solstad *et al*., 2008) (supplementary section: 7). This score takes into account the border coverage by a specific field, its elongation, and its size. Cells having a border score > 0.5 were labelled as border cells.

For later sessions (after object introduction and shift), a new class of neurons pertaining to object representations were observed. Neurons whose spatial information increases by at least 5% and have a spatial information score greater than 0.4 bits/spike (Deshmukh and Knierim, 2011), when an object is introduced into the environment, are classified as object-sensitive cells. To quantify the firing at or around the object location, a Z-score metric was defined by (Tsao, Moser and Moser, 2013). This metric compares the firing rate at the object location to the baseline firing rate away from the object. Neurons that fire near objects (Z-score > 10) are classified as object cells (Deshmukh and Knierim, 2011; Tsao, Moser and Moser, 2013), and other subset of object-sensitive cells fire away from objects (Deshmukh and Knierim, 2011; Høydal *et al*., 2019). Neurons with a Z-score (< 5) before and during object placement but a higher Z-score (>10) after the object is removed are classified as object-trace cells (Tsao, Moser and Moser, 2013).

The hidden layers (P, V, Gr_P, Gr_V, D1, D2, and D3) contain a total of 320 neurons. The above simulation led to a general model that shows a wide range of spatial cells (Fig. 2) when trained with ‘config 111’ (Table 1). These cells were observed in varying proportions (see Table 2) (Fig. 2). We used similar methods to classify cells into specific categories, as done in previous experimental studies. Due to the diverse classification methods inspired by different paradigms, some cells fell into multiple categories. For example, a place cell might also be categorized as an object cell if its firing field remaps with the introduction of an object, or as a trace cell if it disappears with object introduction and reappears at the object’s location when the object is removed. Such cells have been observed in the CA1 and CA3 regions of the hippocampus (Deshmukh and Knierim, 2013).

**Fig 2:**
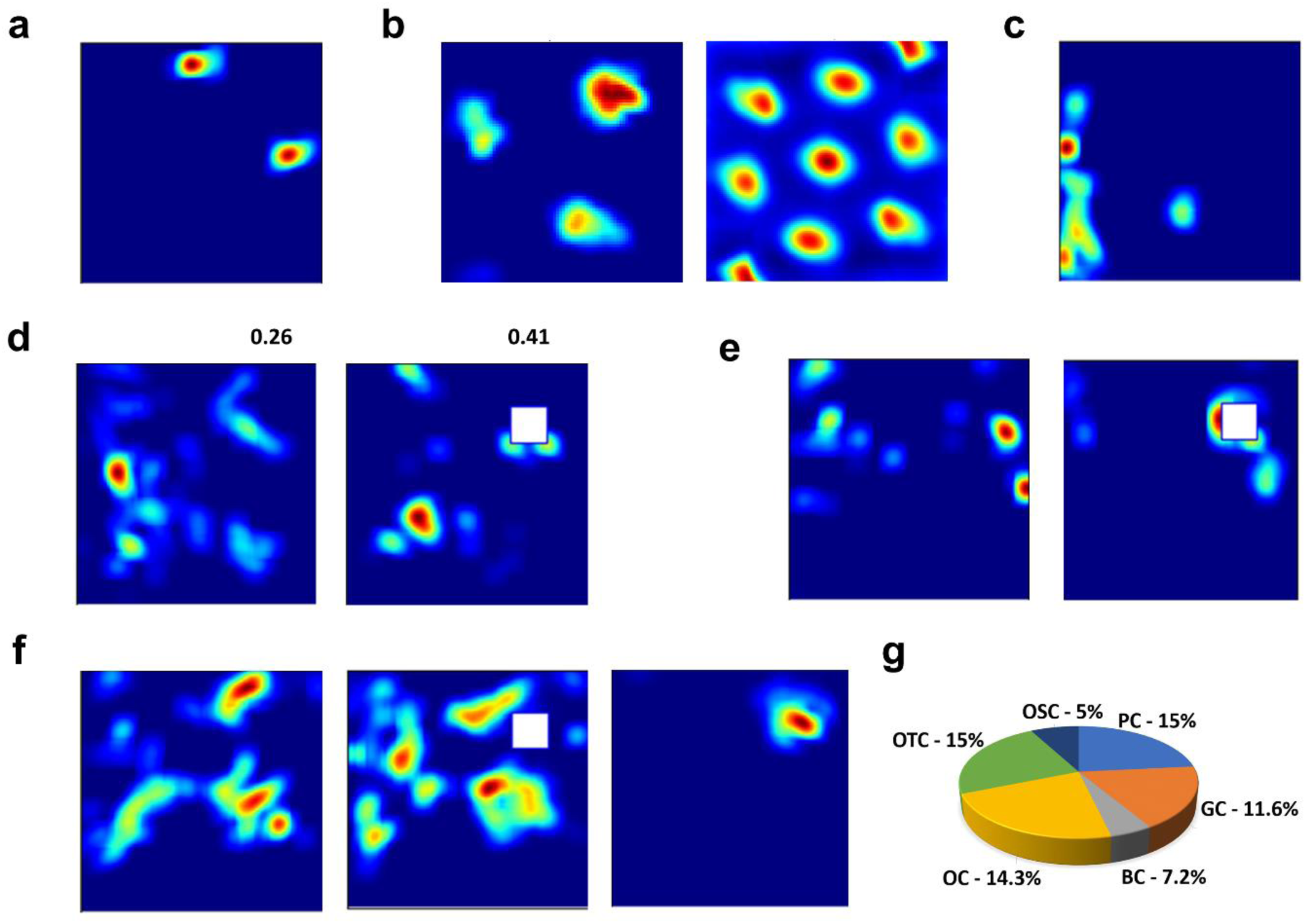
Responses of neurons from hidden layers when the model is trained with ‘config 111’. a-f show examples of different cell types seen in the hidden layers. a) a place cell. b) a grid cell (left) and a spatial 2D auto-correlogram of the grid cell (right). c)a border cell. d) an object-sensitive cell. (Left) response of a neuron without an object in the environment, (right) response of a neuron with an object in the environment. The number on top of each plot denotes the spatial information. e) An object cell. (Left) is the response of the neuron in the absence of an object, and (right) is the response of the neuron after the object is introduced. f) Object-trace cell. (Left) response of the neuron in the absence of an object, (middle) response of the neuron in the presence of an object, and (right) response of the neuron at the location of the object’s previous position after removing the object. g) shows the percentage of each of these cells. PC - place cell, GC - grid cell, BC - border cell, OSC - object sensitive cells, OC - object cell, and OTC – object-trace cell.

**Table 2:**
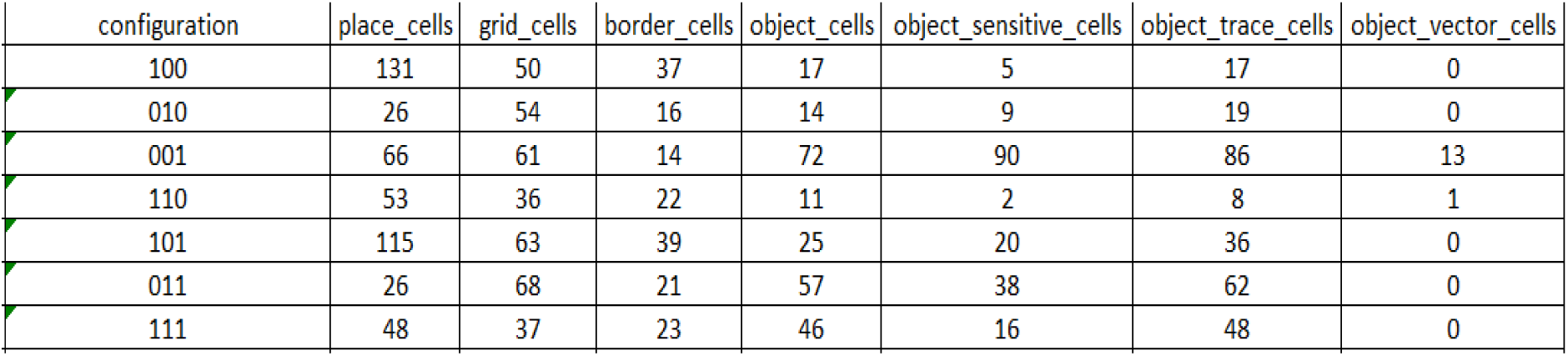
Number of different types of cells for various configurations of learning for position, HD, and reward. (See Table 1 for the definition of the various configurations).

| configuration | place_cells | grid_cells | border_cells | object_cells | object_sensitive_cells | object_trace_cells | object_vector_cells |
| --- | --- | --- | --- | --- | --- | --- | --- |
| 100 | 131 | 50 | 37 | 17 | 5 | 17 | 0 |
| 010 | 26 | 54 | 16 | 14 | 9 | 19 | 0 |
| 001 | 66 | 61 | 14 | 72 | 90 | 86 | 13 |
| 110 | 53 | 36 | 22 | 11 | 2 | 8 | 1 |
| 101 | 115 | 63 | 39 | 25 | 20 | 36 | 0 |
| 011 | 26 | 68 | 21 | 57 | 38 | 62 | 0 |
| 111 | 48 | 37 | 23 | 46 | 16 | 48 | 0 |

This opened a path to explore whether the observed proportion of neurons in the model depend on chosen *config* (Table 1). We found that grid and place cells are present in high proportions under all conditions. Border cells, however, are more common when the model is trained with ‘config 100’. Higher proportions of object-sensitive, object, object vector, and object-trace cells are observed with ‘config 001’. Besides identifying these cells from different model layers, we conducted varied simulations inspired by experimental studies, with the results detailed in the sections below.

### Place cells

We trained a new model with ‘config 100’ as the model showed increased proportions of place cells for these parameters. We classified a total of 131 (∼41%) place cells across all layers based on spatial information (SI) score (mean: 0.35 +- 0.04 bits/spike) and sparsity measure (mean: 0.07 +- 0.02) discussed above. The HGS of these cells were also calculated which came out to be negative, removing the possibility of being a grid cell. Place cells were significant in all layers (P: 44%, GR_V: 35%, GR_P: 56%, D1: 30%, D2: 32%, D3: 60%) except the visual layer (V: 9%) (Fig. 3c). These place cells did not show qualitative differences across layers, as there were no trends in spatial information scores (Fig. 3b, top-right) or sparsity measures (Fig. 3b, bottom-right).

**Figure 3:**
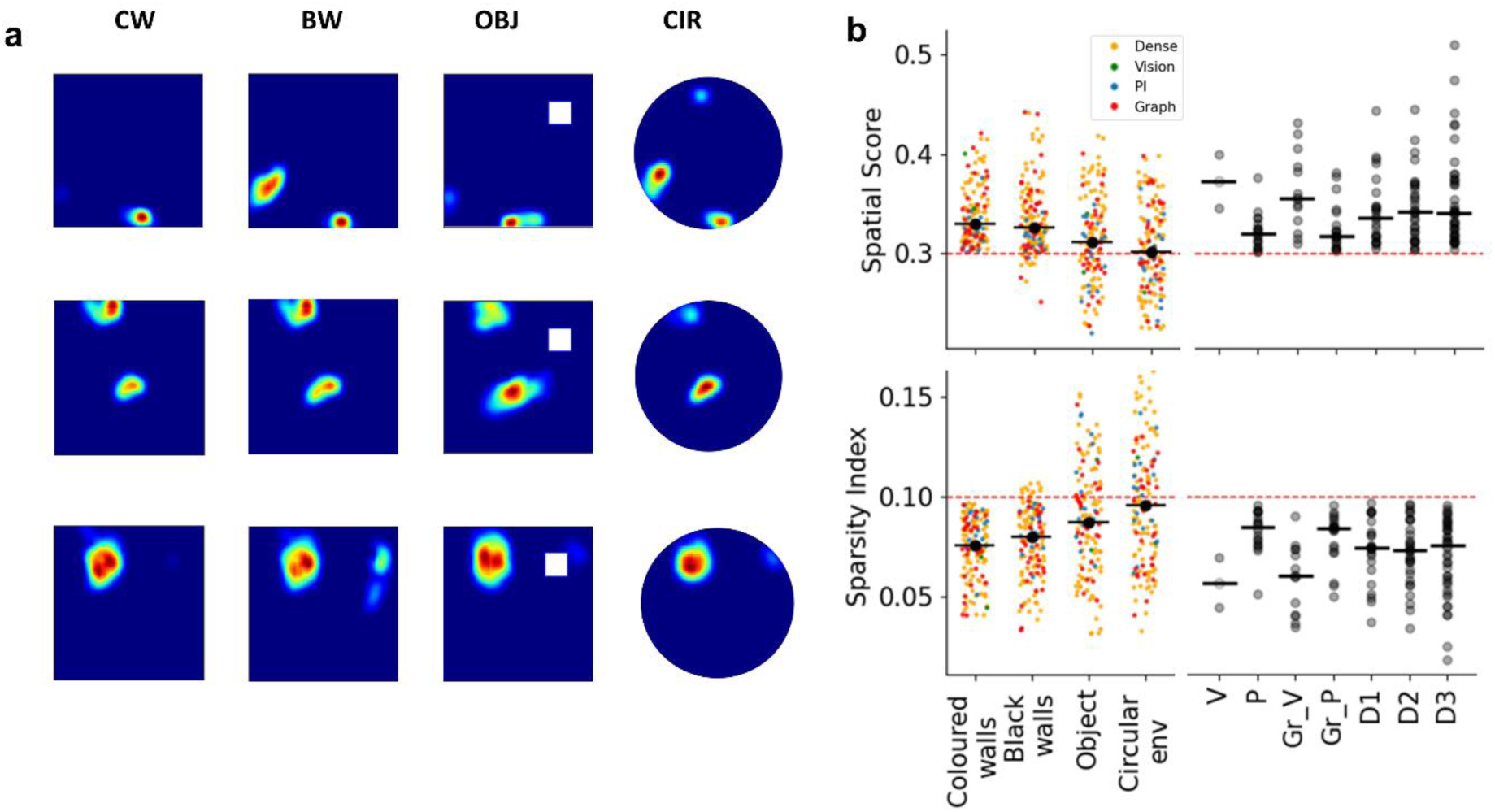
Place cell responses. a) Firing rate map for three different place (top, centre, bottom) cells in three environments. First column: environment with coloured walls and no object. Second column: environment with blackened walls. Third column: an object is introduced to the environment. Object position shown as a white square patch. Fourth column: The environment is changed to a circular environment. b) top: Spatial Information and bottom: sparsity scores for place cells classified in coloured walls environment. left: scores across different environments. Colours denote the layer of the neuron. Vision: V, PI: P, dense: (D1, D2, D3), Graph: (GR_V, GR_P). right: score distribution across layers in coloured walls environment. No clear trend is observed except vision layers have slightly lower spatial specificity. CW: coloured walls, BW: Black walls, OBJ: Object, CIR: Circular environment.

We tested the stability of these place cells by retraining the model under three separate conditions: a) removing distal cues by blackening all walls, b) introducing an object, and c) changing the environment to a circular shape. There was no qualitative change in SI or sparsity for black walls (mean SI: 0.33 +_ 0.06, mean sparsity: 0.08 +_ 0.02), but spatial specificity decreased with object introduction (mean SI: 0.31+ 0.05, mean sparsity: 0.09 +_ 0.03) and circular environment (mean SI: 0.30 +_0.05, mean sparsity: 0.1 +_ 0.03) (Fig. 3b, left). This trend was consistent across all layers (supplementary section: 11, Figure S2). Of the 131 place cells, many retained their spatial properties [black wall: 105, object: 73, circular: 56], while others lost their spatial specificity. The proportion of cells losing spatial specificity was comparable across all layers in all environments (supplementary section: 11, Figure S2: a). We tested place cell remapping or drift by calculating the Euclidean distance between the centroid of the primary field of a place for all three conditions with the general condition (supplementary section: 11, Figure S2: b1, b2, b3, and c). If the field drifted more than 10% of the initial environment size, it was classified as a remapped place cell. Of the cells that retained spatial specificity between environments a fraction of cells remapped [black walls: 14 (∼13%), object: 27 (∼ 37%), circular: 10 (∼18%)].

### Grid cells

We analysed grid cells (Fig. 4a) from the same model used for the place cell in the previous section. Grid cells were classified with a similar method (HGS > 0.1 and 95^th^ percentile from the random shuffling test, supplementary methods). We identified 50 (∼16%) grid cells (mean HGS: 0.46 ± 0.27), distributed irregularly across layers (Fig. 4c). A significant proportion of grid cells were found in the PI layer or layers with direct projections from the PI layer (P: 28%, GR_V: 22%, GR_P: 22%, D1: 30%, D2: 9%, D3: 3%), with none in the Visual layer (V). These grids cells did not show any significant trend in HGS over the visual and dense layers but had slightly higher HGS for PI and PI graph (Fig. 4b, right).

**Figure 4:**
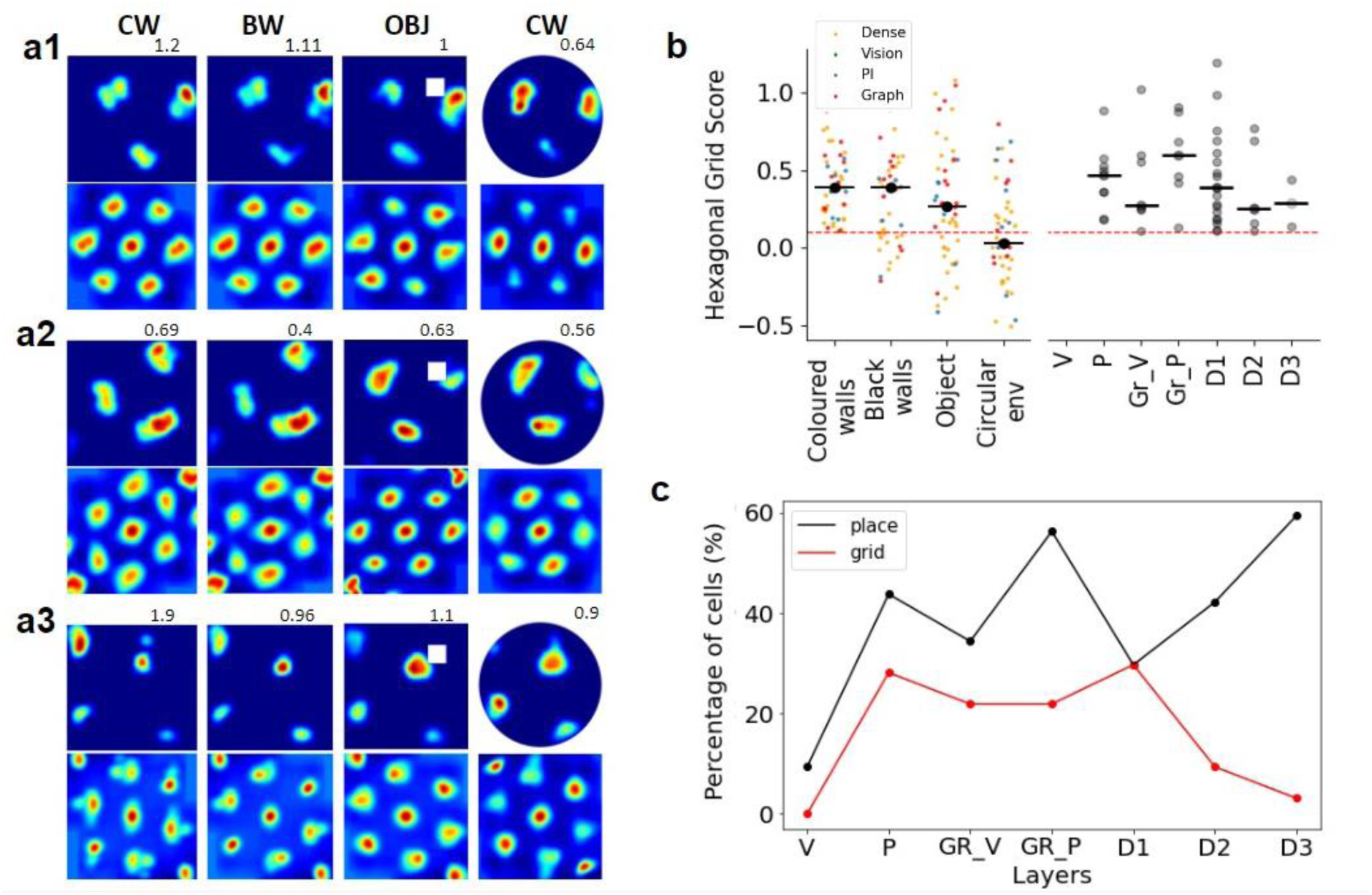
Grid cell responses. a1), a2), a3) firing rate maps (top) and their respective auto-correlogram (bottom) of three grid cells in different environments. First column: grid cell in environment with coloured walls (CW). Second column: response of the same neuron after removing all the visual cues from the environment (black walls). Third column: response of same neuron after introducing the object in the environment (object shown as white patch) . Fourth column: response of same neuron after changing the square environment to a circular environment. b) hexagonal grid scores for grid cells classified in coloured walls environment. left: scores across different environments. Colours denote the layer of the neuron. Vision: V, PI: P, dense: (D1, D2, D3), Graph: (GR_V, GR_P). right: score distribution across layers in coloured walls environment. No clear trend except path integration (PI) layers having slightly higher score than other layers, c) proportion of grid and place cells across different layers. CW: coloured walls, BW: Black walls, OBJ: Object, CW: Circular environment.

To test the stability of these grid cells, we observed their properties in all the three paradigms: black walls, object introduction and circular environment (Fig 4b, left). Of all the 50 grid cells observed, many retained their gridness [black walls: 32 (mean HGS: 0.53 ± 0.25), object: 33 (mean HGS: 0.48 ± 0.28)] but a significant number were lost in a change of environment [circular: 24 (mean HGS: 0.42 ± 0.25)].

### Border representations

Border cells were reported in the subiculum (Hartley and Lever, 2014) and medial entorhinal cortex (Solstad *et al*., 2008), where a family of hippocampal neurons exhibited firing fields elongated along and close to the borders of the environment. A subset of these firing fields was observed to span adjacent borders and were not restricted to a single border.

### Border cells

To increase the relative proportion of the number of border cells, a new model was trained with ‘config 110’ (Table 1). The agent freely traversed a square room (edge length = 1 unit) with coloured walls to provide distal cues but with no landmarks or objects. A border score inspired by (Solstad *et al*., 2008) was employed, and a threshold of > 0.5 border score was applied to filter the border cells. Nearly 27% of all the neurons were classified as border cells (border scores: 0.66 ± 0.09), a quantity higher than observed in experiments (Fig. 5a, top-left). Significant proportion of border cells were observed in visual (V), graph visual (Gr_V) and dense layers (D1, D2, D3) (V: 31%, Gr_V: 31%, D1: 24%, D2: 22%, D3: 28%). Dense layer (D1, D2, D3) neurons had a slightly higher border score than the neurons of the visual layers (V, GR_V) and no border cells were observed in PI layers (P, GR_P) (Fig. 5b).

**Figure 5:**
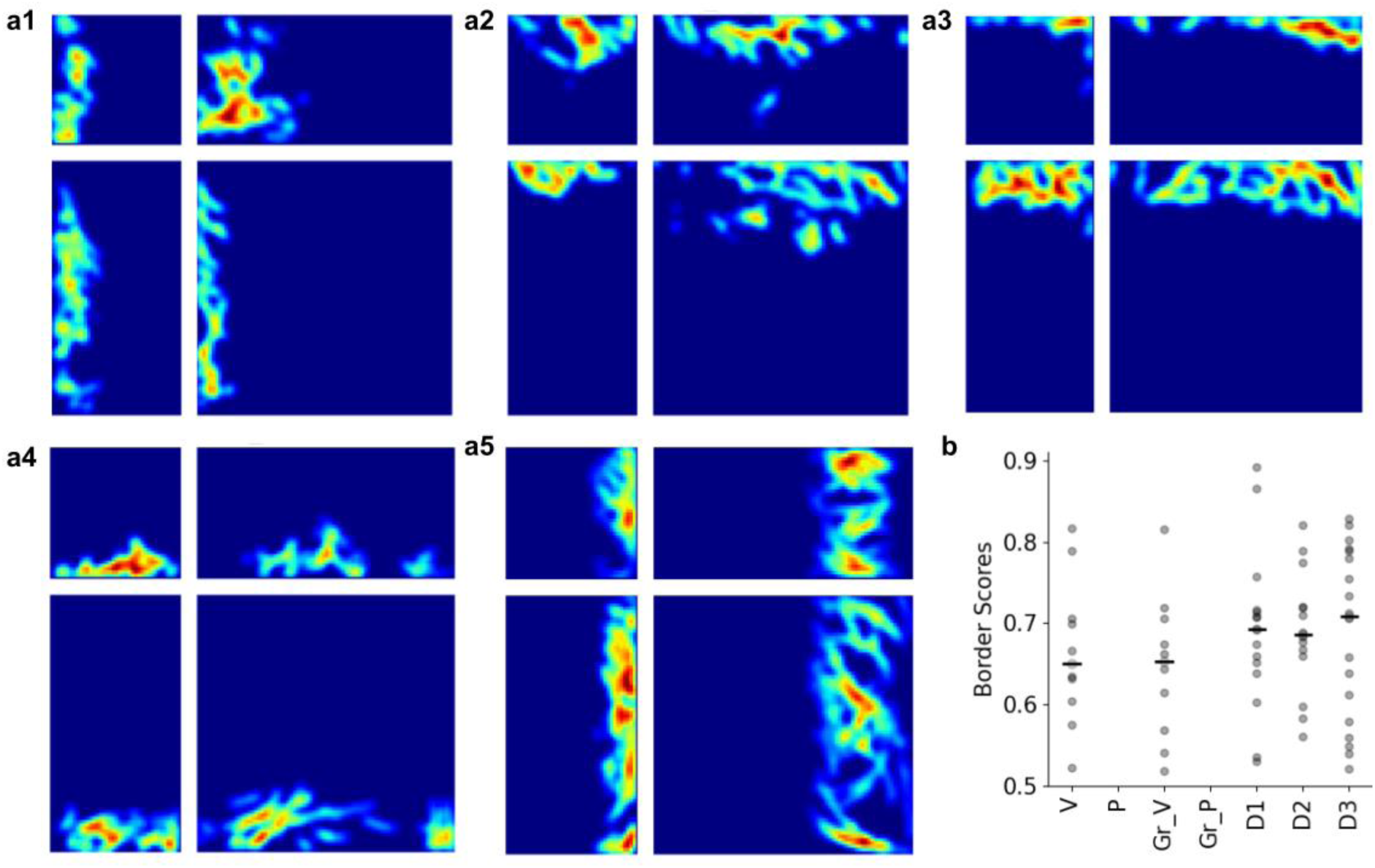
Border cell responses. Five different border cells. a(1-5) are neurons firing show responses of five different border cells in standard size environment (top-left). bottom-right: show responses of same neurons when the size of the environment is doubled. top-right: show responses of same neurons when only the breadth of environment is doubled. bottom-left: show responses of same neurons when only the length of environment is doubled. Neurons for all walls are observed. b) distribution of border scores across layers in standard environment. No border cells are observed in PI and graph-PI (GR_P) layers.

As border and boundary coding is observed to be stable across environments, we extended the environment along its length (Y) (Fig. 5a, bottom-left) and separately along its width (X) (Fig. 5a, top-right) by a factor of 2 and retrained the model for the same. As observed experimentally, the modelled neurons with fields aligned with the extended dimension also exhibited extended firing fields along that border. When the environment was extended about its length and width simultaneously (Fig. 5a, bottom-right), the fields aligned along the length were significantly elongated (t99 = -3.37, p > 0.001; standard env.: 0.753 ± 0.27, extended env.: 0.55 ± 0.26). Similarly, the fields aligned along the width were significantly elongated too (t94 = 0.992, p = 0.32; standard env.: 0.73 ± 0.28, extended env.: 0.68 ± 0.27). Moreover, the fields aligned along unextended borders shifted along with those borders when the environment was expanded (Fig. 5-a4, bottom-left, Fig 5-a5, top-right).

To test the impact of distal cues on border cells, we removed the distal cues from the environment by changing the floor and wall colours to black. As a result, the model learned to predict its current position accurately but could not predict the heading direction. This modification drastically impacted the border cell representations, and only 4 cells of the 320 analysed were classified as border cells. Apart from removing distal cues, we also probed the model to predict position, reward, and heading direction individually and different combinations of all three (Table 1).

### Object Sensitive cells and object cells

Neurons in LEC are more sensitive to the presence of objects in the environment as compared to cells in MEC (Deshmukh and Knierim, 2011; Tsao, Moser and Moser, 2013). These cells were reported as object-sensitive cells. The spatial information of these cells increases with the introduction of objects in the environment.

Although, we have reported object-sensitive cells in our common simulation above. For this study, we found the best results when the model was trained with ‘config 001’ (Table 1). In the first training of the model, the environment with coloured walls and without any object is used. The second step involves retraining the model after introducing the object in the same environment. We classify a neuron as an object-sensitive cell if its spatial information in first session (i.e., without object) is less than 0.4 bits/spike and the difference of spatial information in the second session i.e., after introducing the object in the environment and the first session is more than 0.1 bits/spike. Out of 320 neurons in all the hidden layers, 90 (28.13%) neurons are classified as object-sensitive cells (Fig. 7). As we move from lower to upper layer of the model, a general trend of increase in number of object-sensitive cells is observed (V: 6.25%, P: 12.5%, gr_V: 18.75%, gr_P: 15.63%, D1:31.25%, D2: 37.5%, D3: 45.31%) (Fig. 7b). The mean spatial information of all of these cells before introducing the objects is 0.31 bits/spike whereas it is 0.51 bits/spike after introducing the object in the environment (Fig. 7). In this configuration ‘config 001’ (Table 1), a few of these object-sensitive cells were grid cells (13.33%) in the first training i.e., environment without object and showed object-sensitive-like behaviour after introducing the object into the environment. Similarly, 20% of object-sensitive cells were place cells. Among the neurons that meet the criteria of object-sensitive cells, 50% of neurons fire near the object with z-score >= 10 and the remaining 50% fire away from the object with z-score < 10.

**Figure 7:**
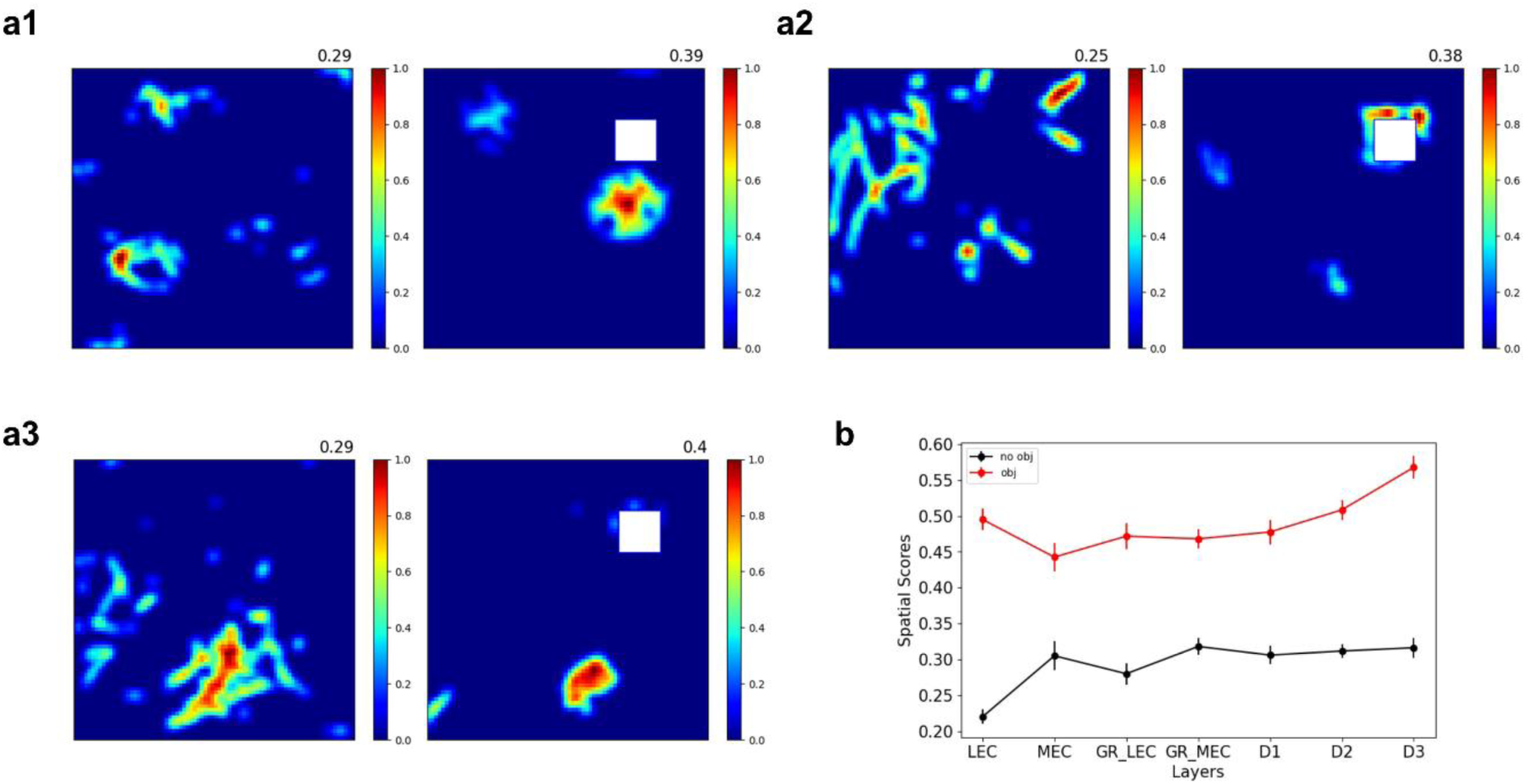
Objective sensitive cells. a1, a2 and, a3 depict the responses of three object-sensitive cells. a1) (left) firing rate map of neuron when object is not present in the environment and (right) firing rate map of same neuron after introducing the object in the environment. The number on top shows the spatial information in each case. a2 and a3 are similar to a1. b) shows the average spatial information of object-sensitive cells in each layer in the environment without object and with object.

This classification of object-sensitive cells into near-object firing cells and away-from-object firing cells was done by (Deshmukh and Knierim, 2011). Although, in later extensive studies, neurons that fired near the object were named object cells (Tsao, Moser and Moser, 2013). In our simulation experiments, we found 72 (22.5%) object cells that fired near the object upon introducing the object to the environment. The model was retrained after shifting the object to a new location, and 36 (50%) neurons shifted their firing along with the object. The remaining neurons either fired at the previous location (called object-trace cells: 5.6%) or their firing remapped (94.4%). The model was again retrained with a differently shaped object (a spherical object), and from the 72 object cells 86.1% neurons did not lose their object cell property while the firing fields of the remaining neurons remapped (Fig. 8).

**Figure 8:**
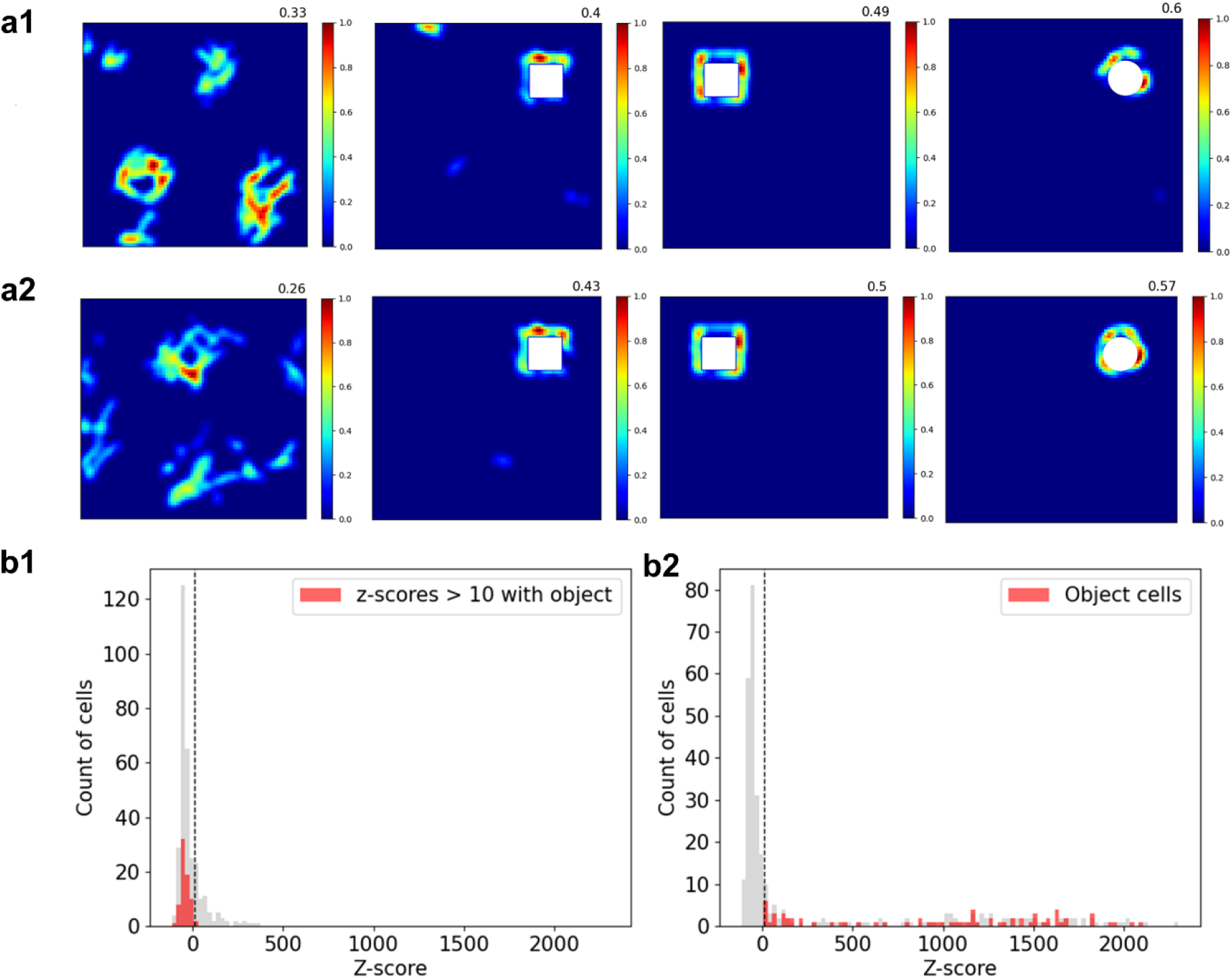
a1 and a2 are two different object cells showing response in four sessions. In a1 first column is the firing rate map of neuron when the object is not present. Second column is the firing rate map of neuron when a cubical object is introduced in the environment. Third column is the firing rate map of the same neuron when the object is shifted to a new location and fourth column is the firing rate map of the neuron when the cubical object is replaced by a spherical object. a2 is same for another neuron. b1 shows the z score of object cells before introducing the object and b2 shows the z score of the object cells after introducing the object in the environment.

### Object vector cells

Neurons that form a vectorial relationship with the object, i.e., those that fire at a particular distance and direction from the object, are called object vector cells (Deshmukh and Knierim, 2013; Høydal *et al*., 2019). In this study, we found the best results when the model was trained with ‘config 001’ (Table 1). This study is done on a total of 8 training sessions. In the first session, the model is trained without an object. In the subsequent sessions, the model is retrained in a sequential manner. In the second session, an object is introduced in the environment and a firing field emerges and gets associated with the object. In the third session, the object is shifted to a new location, and the firing field also shifts, maintaining the same vectorial relationship with the object. In the fourth session, we replace the cubical object with a spherical object at the second session’s object location. In the fifth session, we replaced the spherical object with the cylindrical object. The firing field still maintains the vectorial relationship with the object. In the 6th, 7th, and 8th sessions, the cubical object from session two is extended gradually, and each time the model is retrained. Neurons B, C, and D maintain the vectorial relationship while neuron A’s firing field remaps. To quantify the vectorial relationship, an object vector score is calculated between two sessions (see Supplementary section 1). The correlation of the distance and direction of each spike is calculated from the centre of the object between different sessions. A neuron is classified as an object vector cell if the score is less than 0.4 between session 1 and session 2, i.e., between no object and introduction of a cubical object, and more than 0.4 between session 2 and every other subsequent session. Out of all the neurons in hidden layers, we found 13 (V: 3.1%, D1: 7.8%, D2: 4.7%, D3: 4.7%) neurons that show object vector-like responses. Among these 13 neurons, one of the neurons maintained the vector relationship only up to session 3. Out of the remaining 12 neurons, 3 maintained their vector relationship up till session 5, and the remaining 9 neurons maintained the vector relationship till session 8 (Fig. 9).

**Figure 9:**
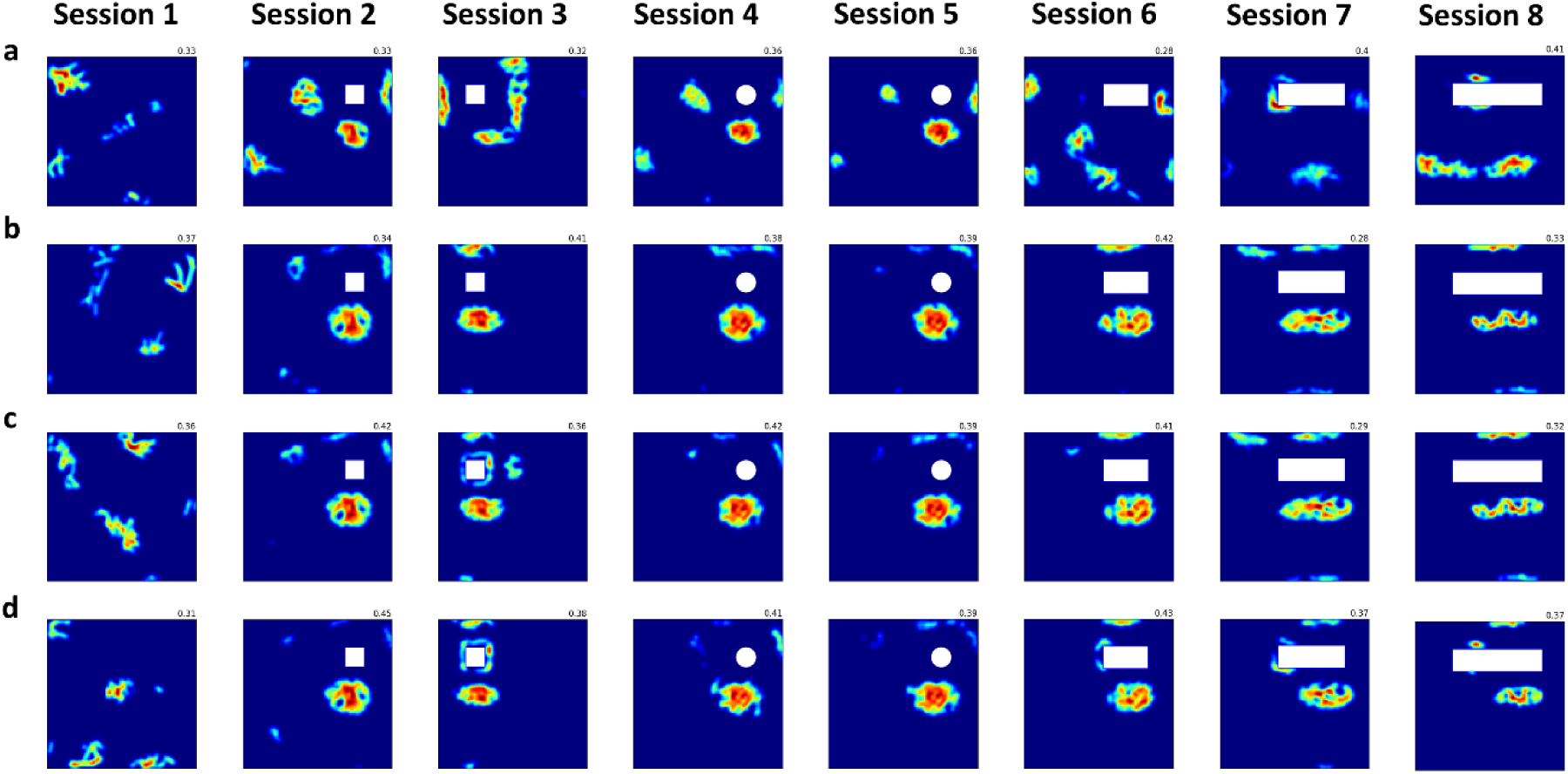
Object vector cell responses. The four rows show the response of 4 different object vector cells. a) is the firing rate map of a neuron across 8 sessions. Session 1 shows the response of neurons when the object is not present. Session 2 shows the emergence of a firing field associated with the object when the object is introduced. Session 3 shows the shifting of the firing field with the shift of the object, maintaining the vectorial relationship with the object. Session 4 shows the firing field after changing the shape of the object to the sphere. Session 5 is the same as Session 4 for cylindrical object. Sessions 6, 7, and 8 show the firing response as the object is extended gradually. Note that a’s firing gets distorted as the object is extended, whereas neurons b, c, and d maintain the vector relationship with the extended object.

**Figure 10:**
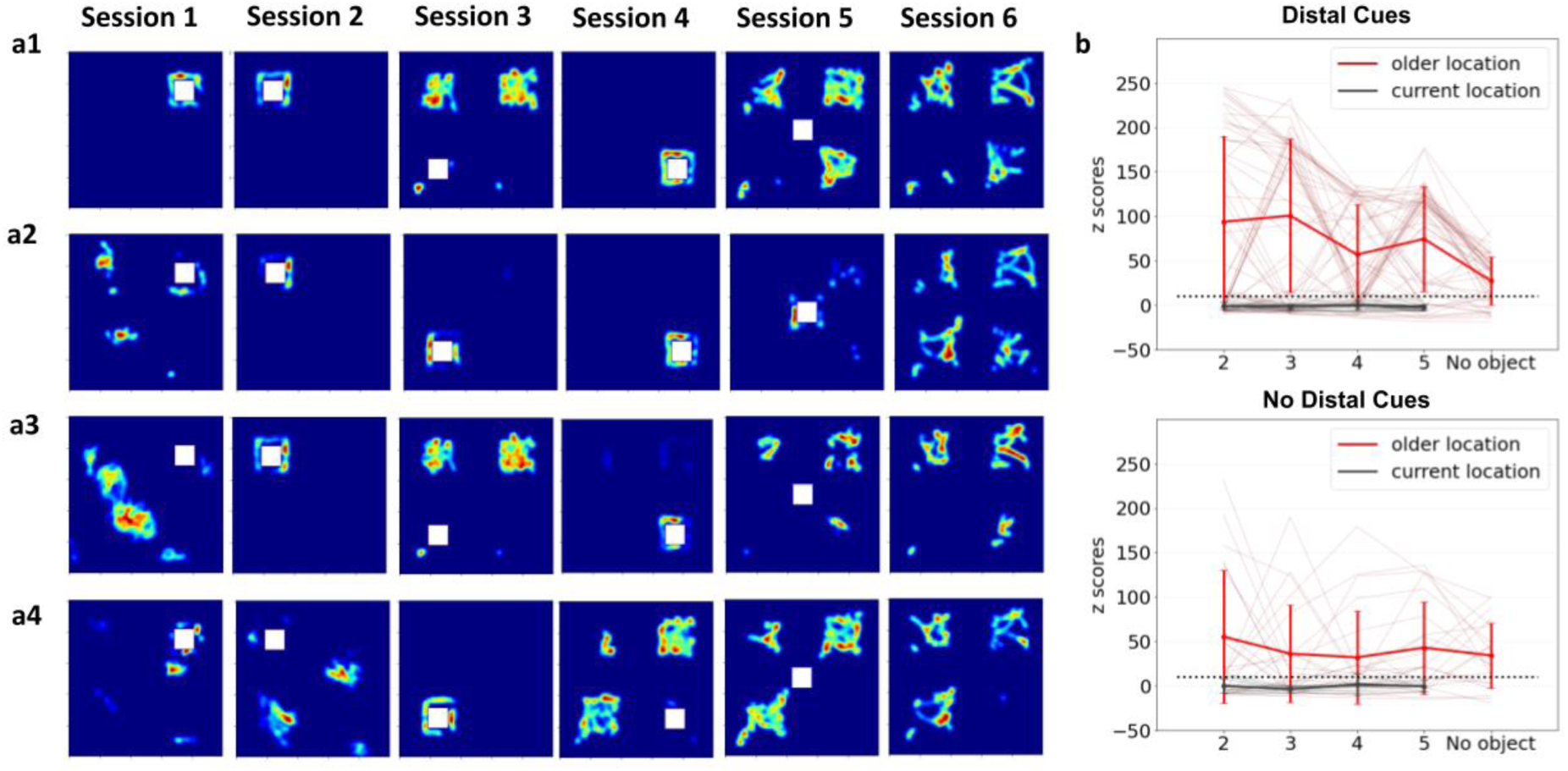
Object-trace cell responses. a(1-4) depicts the 6 sessions of four object-trace cells from the hidden layers of the model. a1 shows the firing fields near the object while training the model in session 1. Session 2: the firing response of the same neuron after shifting the location of the object. The firing field emerges near the current location of object. Session 3: the response after training the model in third session. The firing fields at the last two location emerges while little response is seen near the third location of the object. Session 4: the firing field emerges near the object although it disappeared near previous locations. Session 5: retraining of the model when the object is present at the centre of the environment. Session 6: the firing field of the neuron after retraining the model again in ‘without object’ condition. The neuron is able to remember three previous locations of the object properly. Similarly, the response of neuron a(2-4) can also be observed during different sessions. b) Z scores of neurons with coloured walls (top) and black walls (bottom). Thick lines represent mean and thin lines represent z-score of individual neurons from session 2-6.

### Object-trace cells

In this study, we investigated the properties of object-trace cells, which encode prior object locations in an environment reported by (Deshmukh and Knierim, 2013; Tsao, Moser and Moser, 2013). The configuration used for object-trace cells is ‘config 001’ (Table 1), across 6 training sessions designed to emulate the aforementioned studies (Fig. 8a). In the 1^st^ session, the agent freely traverses along a random trajectory in a square environment with colored walls and a black floor. For the 2^nd^ session, a cubical object is introduced to the environment, and the model is retrained. Furthermore, the object is removed, and the model is retrained. This training procedure is simulated to replicate the experimental method used by (Tsao, Moser and Moser, 2013) to classify object-trace cells. We assessed neuronal response to the object using a Z-score metric comparing firing rates near the older object location to other areas.

The object was sequentially relocated in sessions three to five, and the model was retrained at each new position. The final session involved retraining with no object present, yet trace firing persisted. The same protocol was repeated in an environment with black walls to investigate the impact of distal cues and context on memory retention. A significant decrease occurred in the percentage of trace cells maintaining their firing patterns when retrained with new object placements in a black-walled environment [coloured walls: 51 (16%), black walls: 21 (7%)] (Fig. 8b). This can be posited as a catastrophic forgetting problem in a continual learning paradigm where older representations are erased when retrained with novel data. Nevertheless, this needs to be addressed to capture the object-trace cell properties. To mitigate this, we applied Elastic Weight Consolidation (EWC) (Kirkpatrick et al., 2017), which uses regularization to maintain critical parameter stability during new task training. While EWC did not affect trace cell proportions in the colored wall environment, it significantly enhanced them in the black wall setting [supplementary section: 12, Figure S3], demonstrating the importance of contextual cues in object memory tasks.

## DISCUSSION

Organisms integrate information obtained from multiple sensory modalities to navigate in the three-dimensional world and visual landmarks serve as references for such navigation. Although there are a large number of modelling studies that model place and grid cells individually (Solstad, Moser and Einevoll, 2006; Hasselmo, Giocomo and Zilli, 2007; Giocomo and Hasselmo, 2008; Burgess and O’Keefe, 2011), there is a need for a comprehensive model that integrates all spatial and object-responsive cells.

In this study, we developed a comprehensive deep learning model with two input streams: Path Integration and vision analogous to the brain’s ‘where’ and ‘what’ pathways (Deshmukh and Knierim, 2011; Save and Sargolini, 2017) producing various hippocampal representations in the hidden layers of the model. The model’s key element is the Graph Convolution Network (GCN) which performs sensory integration of locomotion (PI) and vision. The GCN layer establishes lateral connectivity between sensory modalities, similar to those in the LEC and MEC regions of the Entorhinal Cortex (Deshmukh and Knierim, 2011; Save and Sargolini, 2017) that gives it memory capability. Unlike previous models, the GCN does not assume any input graph structure, as trajectories are randomly generated so as to uniformly cover the 2D space. Furthermore, pertaining to GCN the model does not assume any graph structure in the input contrary to previous models (Whittington *et al*., 2020) as the trajectories are stochastically generated with uniform distribution across 2D space. As the individual sensory modality is posited as a graph node, the lateral connectivity between nodes and impact of each input on representations can be investigated separately. Additionally, GCN provides a modular framework for sensory integration such that each sensory modality can be modelled independently before graph convolution.

While the model is trained to predict current position, heading direction, and reward, the specific neuron types emerge naturally in the hidden layers. The model shows emergence of place, grid, border, boundary-vector, object-sensitive, object, object-vector, and object-trace cells. Although, the relative proportions of these cells depend on the training with different configurations as discussed in Table 1. A mixed proportions of these responses across hidden layers of the model is observed. However, there is a preference of layers for particular neuron types. Object vector cells predominantly appear in higher layers after the GCN, as both PI and vision are needed to associate the object with the agent’s location and direction. Similarly, higher proportions of object-trace, object sensitive and object cells are observed in post GCN layers.

The observed proportions of neural representations depend significantly on the configuration (Table 1) of the ground truth. For example, training with ‘config 100’ results in more border cells. On the contrary, ‘config 001’ leads to higher proportions of object-related firing. In such cases, the hidden layer neurons predominantly act as object-sensitive, object-vector, and object-trace cells. Similarly, place and grid cells invariably emerged for all configurations, but their proportions increased in ‘config 110’. As we perform an exhaustive search through all possible configurations and investigate how the different configurations impact hidden layer responses to study these cells independently and infer their functional roles in navigation.

We first trained a basic model with ‘config 111’ and reported the spatial and object-related neurons. Given the abundance of place and grid cells, we further analysed them under different environmental conditions. Introducing an object and retraining the model revealed that some neurons’ firing locations remained unchanged, while others remapped, reminiscent of the experimental studies (Chen *et al*., 2013) for motion-based, vision-based, and conjunctive place cells. We further analysed the place and grid cells after removing visual inputs by making the environment’s walls and floor black; similar trends were observed, with many neurons maintaining their grid or place cell properties.

Next, we analysed border cells in our model. The highest proportion of border cells was observed when the agent is trained with ‘config 110’ with a necessary condition of presence of position in the configuration. The border cells also highly depend on the visual input.

The object-based representations including object sensitive cells, object cells, object-vector cells and object-trace cells were dominantly present when the model was trained with ‘config 001’. This indicates that animals associate objects with reward while navigating as suggested before (Davidow *et al*., 2016; Loh *et al*., 2016). Our model replicated experimental findings (Deshmukh and Knierim, 2011)), showing higher spatial information in the presence of objects than in their absence. We analysed neurons based on their firing locations and found many firing near the object, consistent with experimental studies (Deshmukh and Knierim, 2011; Tsao, Moser and Moser, 2013). Many neurons maintained their object cell properties even when the object was moved or changed, while some did not. Neurons that lost this property when the object shifted sometimes fired at the previous location, indicating they encode a memory of the object’s previous location. To address continual learning with multiple object shifts, we employed Elastic Weight Consolidation (EWC) in the model. This approach maintains critical weights from previous tasks, increasing memory cell proportions and network capacity. The model could also encode the agent’s position and direction from the object through OVC. The OVCs were again tested on varying environmental conditions that matched the experimental study (Høydal *et al*., 2019).

The model successfully demonstrated the emergence of experimentally observed spatial cells across multiple training paradigms. However, deviations from published literature were noted. Sometimes, the same cell could be classified into multiple categories: for instance, a place cell could remap with object introduction to exhibit object cell or trace properties. The hippocampus integrates information from multiple sources, leading to single cells with contextual firing fields exhibiting both object and memory features, as seen in the CA1 region (Deshmukh and Knierim, 2013). Additionally, some modelled grid cells lost their grid properties when an object was introduced or the environment changed. This requires further investigation, but recent studies (Boccara *et al*., 2019) have shown that reward impacts grid cell properties, which is also observed in our model. Additionally, when visual input was removed, the number of border cells in the model drastically decreased. This indicates that distal cues are essential for border cell emergence, contrasting with findings by (van Wijngaarden, Babl and Ito, 2020) where border cells were observed in the dark and their firing was invariant to distal cue rotation. This issue might be resolved by adding a border proximal input, such as haptic input, to the GCN layer, in addition to PI and vision. Additionally, when the visual input was removed, the number of border cells drastically decreased in the model. Hence, visual cues are essential for the emergence of border cells, which contrasts with (van Wijngaarden, Babl and Ito, 2020) where, the border cells are observed in dark the and the border cell’s firing was invariant with rotation in distal cues. This can be further solved by adding a border proximal input such as the haptic input to the GCN layer, apart from PI and vision.

Our model is fashioned after deep neural networks and may perhaps not possess biophysical realism. However, it captures the hippocampal data flow, with PI input capturing the “where” pathway and visual input capturing the “what” pathway (Deshmukh and Knierim, 2011), both merging in the hippocampus, similar to the GCN in our model. Currently, input to the PI pipeline is modelled as a sinusoidal function of the position, after eliminating temporal information by time-averaging (supplementary section: 9). PI actually integrates velocity information to estimate position. A land animal gets its self-motion information from the proprioceptive feedback coming from its locomotion. Therefore, in future studies, one can go further and actually model limb oscillation (Soman, Muralidharan and Chakravarthy, 2018) and present it as input to the PI pipeline. Also, other sensory modalities can be added such as auditory and somatosensory inputs along with vision and PI and can further be probed for more complex environmental paradigms. The current model only describes the emergence of spatial and object related cells in the hippocampal formation. It describes spatial representations of cells and not navigation. Future work entails expanding the model to be a complete Reinforcement Learning agent that navigates different natural environments and investigate the neural responses.

## REFERENCES

Aziz, A. et al. (2022) ‘An integrated deep learning-based model of spatial cells that combines self-motion with sensory information’, *Hippocampus*. John Wiley & Sons, Ltd. doi: 10.1002/HIPO.23461.

Bjerknes, T. L., Moser, E. I. and Moser, M. B. (2014) ‘Representation of geometric borders in the developing rat’, Neuron. Cell Press, 82(1), pp. 71–78. doi: 10.1016/J.NEURON.2014.02.014.

Boccara, C. N. et al. (2019) ‘The entorhinal cognitive map is attracted to goals’, Science, 363(6434), pp. 1443–1447. doi: 10.1126/science.aav4837.

Burak, Y. and Fiete, I. R. (2009) ‘Accurate Path Integration in Continuous Attractor Network Models of Grid Cells’, PLoS Comput Biol, 5(2), p. 1000291. doi: 10.1371/journal.pcbi.1000291.

Burgess, N. and O’Keefe, J. (2011) ‘Models of place and grid cell firing and theta rhythmicity’, Current Opinion in Neurobiology, pp. 734–744. doi: 10.1016/j.conb.2011.07.002.

Chan, E. et al. (2012) ‘From objects to landmarks: The function of visual location information in spatial navigation’, *Frontiers in Psychology*. Frontiers, 3(AUG), p. 25389. doi: 10.3389/FPSYG.2012.00304/BIBTEX.

Chen, G. et al. (2013) ‘How vision and movement combine in the hippocampal place code’, Proceedings of the National Academy of Sciences of the United States of America, 110(1), pp. 378–383. doi: 10.1073/pnas.1215834110.

Clark, R. E. et al. (2002) ‘Anterograde Amnesia and Temporally Graded Retrograde Amnesia for a Nonspatial Memory Task after Lesions of Hippocampus and Subiculum’, Journal of Neuroscience. Society for Neuroscience, 22(11), pp. 4663–4669. doi: 10.1523/JNEUROSCI.22-11-04663.2002.

Cueva, C. J. and Wei, X.-X. (2018) ‘Emergence of grid-like representations by training recurrent neural networks to perform spatial localization’, arXiv. arXiv.

Davidow, J. Y. et al. (2016) ‘An Upside to Reward Sensitivity: The Hippocampus Supports Enhanced Reinforcement Learning in Adolescence’, Neuron. Cell Press, 92(1), pp. 93–99. doi: 10.1016/J.NEURON.2016.08.031.

Deshmukh, S. S. and Knierim, J. J. (2011) ‘Representation of non-spatial and spatial information in the lateral entorhinal cortex’, Frontiers in Behavioral Neuroscience. Frontiers Media SA, 5(OCTOBER). doi: 10.3389/fnbeh.2011.00069.

Deshmukh, S. S. and Knierim, J. J. (2013) ‘Influence of local objects on hippocampal representations: Landmark vectors and memory’, *Hippocampus*. John Wiley & Sons, Ltd, 23(4), pp. 253–267. doi: 10.1002/hipo.22101.

Fyhn, M. et al. (2004) ‘Spatial representation in the entorhinal cortex’, Science. American Association for the Advancement of Science, 305(5688), pp. 1258–1264. doi: 10.1126/SCIENCE.1099901/SUPPL_FILE/FYNH.SOM.PDF.

Gilbert, P. E. and Brushfield, A. M. (2009) ‘The role of the CA3 hippocampal subregion in spatial memory: A process oriented behavioral assessment’, Progress in Neuro-Psychopharmacology and Biological Psychiatry, 33(5), pp. 774–781. doi: 10.1016/j.pnpbp.2009.03.037.

Giocomo, L. M. and Hasselmo, M. E. (2008) ‘Computation by oscillations: Implications of experimental data for theoretical models of grid cells’, Hippocampus, 18(12), pp. 1186–1199. doi: 10.1002/HIPO.20501.

Girardeau, G. et al. (2009) ‘Selective suppression of hippocampal ripples impairs spatial memory’, Nature Neuroscience, 12(10), pp. 1222–1223. doi: 10.1038/nn.2384.

Gothard, K. M. et al. (1996) ‘Binding of Hippocampal CA1 Neural Activity to Multiple Reference Frames in a Landmark-Based Navigation Task’, The Journal of Neuroscience, 16(2), pp. 823–835.

Griffin, A., … H. E.-J. of and 2007, undefined (2007) ‘Spatial representations of hippocampal CA1 neurons are modulated by behavioral context in a hippocampus-dependent memory task’, Soc Neuroscience, 27(9), pp. 2416–2423. doi: 10.1523/JNEUROSCI.4083-06.2007.

Hafting, T. et al. (2005) ‘Microstructure of a spatial map in the entorhinal cortex’, Nature. Nature Publishing Group, 436(7052), pp. 801–806. doi: 10.1038/nature03721.

Hartley, T. and Lever, C. (2014) ‘Know your limits: The role of boundaries in the development of spatial representation’, Neuron. Cell Press, 82(1), pp. 1–3. doi: 10.1016/J.NEURON.2014.03.017.

Hasselmo, M. E., Giocomo, L. M. and Zilli, E. A. (2007) ‘Grid cell firing may arise from interference of theta frequency membrane potential oscillations in single neurons’, Hippocampus, 17(12), pp. 1252–1271. doi: 10.1002/HIPO.20374.

Høydal, Ø. A. et al. (2019) ‘Object-vector coding in the medial entorhinal cortex’, Nature. Nature Publishing Group, 568(7752), pp. 400–404. doi: 10.1038/s41586-019-1077-7.

Kipf, T. N. and Welling, M. (2016) ‘Semi-Supervised Classification with Graph Convolutional Networks’. Available at: http://arxiv.org/abs/1609.02907.

Loh, E. et al. (2016) ‘Context-specific activation of hippocampus and SN/VTA by reward is related to enhanced long-term memory for embedded objects’, Neurobiology of Learning and Memory. Elsevier, 134(Pt A), p. 65. doi: 10.1016/J.NLM.2015.11.018.

O’ Keefe, J. and Burgess, N. (1996) ‘Geometric determinants of the place fields of hippocampal neurons’, Nature, 381(6581), pp. 425–428. doi: 10.1038/381425A0.

O’Keefe, J. and Dostrovsky, J. (1971) ‘The hippocampus as a spatial map. Preliminary evidence from unit activity in the freely-moving rat’, Brain Research, 34(1), pp. 171–175. doi: 10.1016/0006-8993(71)90358-1.

Sargolini, F. et al. (2006) ‘Conjunctive representation of position, direction, and velocity in entorhinal cortex’, Science, 312(5774), pp. 758–762. doi: 10.1126/SCIENCE.1125572.

Save, E. and Sargolini, F. (2017) ‘Disentangling the role of the MEC and LEC in the processing of spatial and non-spatial information: Contribution of lesion studies’, Frontiers in Systems Neuroscience. Frontiers Media S.A., 11, p. 81. doi: 10.3389/FNSYS.2017.00081/BIBTEX.

Scarselli, F. et al. (2009) ‘The Graph Neural Network Model’, IEEE Transactions on Neural Networks, 20(1), pp. 61–80. doi: 10.1109/TNN.2008.2005605.

Skaggs, W. E. et al. (1994) ‘A Model of the Neural Basis of the Rat’s Sense of Direction’, Advances in Neural Information Processing Systems, 7.

Solstad, T. et al. (2008) ‘Representation of Geometric Borders in the Entorhinal Cortex’, Science, 322(5909), pp. 1865–1868. doi: 10.1126/science.1166466.

Solstad, T., Moser, E. I. and Einevoll, G. T. (2006) ‘From grid cells to place cells: A mathematical model’, Hippocampus, 16(12), pp. 1026–1031. doi: 10.1002/hipo.20244.

Soman, K., Muralidharan, V. and Chakravarthy, S. (2018) ‘A unified hierarchical oscillatory network model of head direction cells, spatially periodic cells, and place cells’, *European Journal of Neuroscience*. John Wiley & Sons, Ltd, 47(10), pp. 1266–1281. doi: 10.1111/EJN.13918.

Tsao, A., Moser, M. B. and Moser, E. I. (2013) ‘Traces of experience in the lateral entorhinal cortex’, *Current biology : CB*. Curr Biol, 23(5), pp. 399–405. doi: 10.1016/J.CUB.2013.01.036.

Whittington, J. C. R. et al. (2020) ‘The Tolman-Eichenbaum Machine: Unifying Space and Relational Memory through Generalization in the Hippocampal Formation’, Cell. Elsevier, 183(5), p. 1249. doi: 10.1016/J.CELL.2020.10.024.

van Wijngaarden, J. B. G., Babl, S. S. and Ito, H. T. (2020) ‘Entorhinal-retrosplenial circuits for allocentric-egocentric transformation of boundary coding’, eLife. eLife Sciences Publications Ltd, 9, pp. 1–25. doi: 10.7554/ELIFE.59816.

Young, B. J. et al. (1997) ‘Memory Representation within the Parahippocampal Region’, Journal of Neuroscience. Society for Neuroscience, 17(13), pp. 5183–5195. doi: 10.1523/JNEUROSCI.17-13-05183.1997.

