## Supplementary Information for "A unified model of hippocampal spatial and object cells involving bidirectionally coupled Lateral and Medial Entorhinal Cortical layers"

**Supplementary Materials**

1. **Object Vector Score (OVS):** To calculate OVS, we calculate the distance and direction of each spike’s position from the centre of the object. For session one i.e., the session without any object, we keep the centre of the object similar to that of session two. A firing rate map is plotted in a distance vs direction plot, and correlation is calculated. For a neuron to be classified as OVC, the correlation between session 1 and session 2 should be less than 0.4 in order to remove the possibility of the neuron being a place cell, and the correlation between session 2 and remaining sessions should be more than 0.4 (Fig. S1).

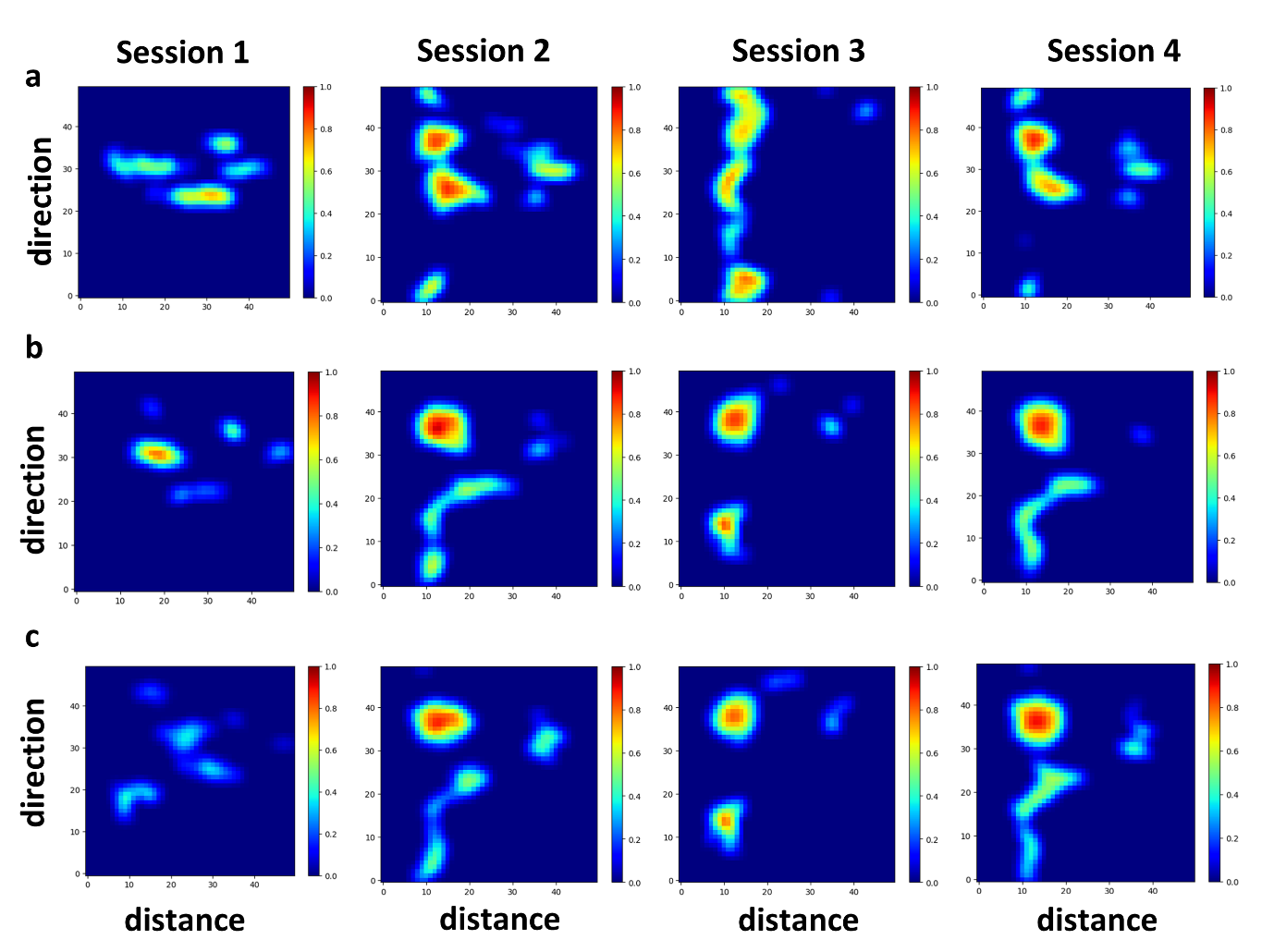

Figure S1: Firing rate map of spikes in a distance vs direction plot for session 1 to session 4 for neurons a, b, and, c corresponding to neurons in Fig. 9.

2. **Elastic Weight Consolidation (EWC):** Elastic Weight Consolidation (EWC) means regularizing the loss in model with new task in such a way that the weights important to previous task does not deviate much from the weights obtained after training the model with previous task (Kirkpatrick *et al.*, 2017). The loss function L in the new task that is minimized using EWC is as follows:

$$L\left( \theta\right)= L_{{session}_{n}}+ \sum_{i} \frac{\lambda}{2}F_{ij}\left( \theta_{i}- \theta_{{session}_{n-1}, i}^{*} \right)^{2}$$

where, $L_{{session}_{n}}$ is the loss for current session, $\lambda$ is the weightage given to older weights as compared to new weights and, $i$ iterates over all parameters, $\theta$ are the current parameters and $\theta$^*^ are parameters from previous training session.

$F_{ij}$ is the Fisher matrix computed by the outer product of gradient of log-likelihood of loss with respect to all parameters from previous session. It is given by:

$$F_{ij}= E_{\left( x,y \right)\sim D}\left[ \frac{\partial log P(y|x,\theta)}{\partial\theta_{i}} . \frac{\partial log P(y|x,\theta)}{\partial\theta_{j}} \right]$$

The diagonal elements of the Fisher matrix represent the curvature of the log likelihood with respect to each parameter. Therefore, in EWC the diagonal elements are used for regularization to prevent significant changes in the parameters when learning new tasks.

3. **Firing Rate Map:** Rate maps for each neuron in each environment was constructed by dividing the environment into a grid of (80*80) spatial bins. We then calculate the firing rate for each bin by dividing the number of spikes recorded in that bin by the number of trajectory points sampled in that bin. To ensure the data's reliability, we smooth the firing rates using a Gaussian kernel with a window size of 5*5. This process results in a firing rate map, which visually represents the spatial locations where a neuron is active.
4. **Sparsity**

Sparsity is determined by taking the square of average firing rate and dividing it by the weighted sum of the squared firing rates. This method ensures that the calculated sparsity accurately reflects the neuron's selectivity across different positions. Mathematically, sparsity *S* is given by:

$$S= \frac{\sum_{i} {(p}_{i}* r_{i}^{2})}{\sum_{i} {(p_{i}* r_{i})}^{2}}$$

where *p_i_* are the elements of the normalized position probability in the i_th_ bin of occupancy map and *r_i_* is the firing rate for i_th_ bin in firing rate map for the neuron.

5. **Spatial Information**

The spatial information index measures the amount of information, in bits, that a single spike conveys about an animal’s location, effectively assessing the mutual information between an animal’s position and a cell’s firing rate. The spatial information content of a cell's discharge is calculated using the following formula:

$$Information Content= \sum P_{i}*\left( \frac{R_{i}}{R} \right)*\log_{2} \left( \frac{R_{i}}{R} \right)$$

where *i* represents the bin number, *P_i_* is the probability of the animal occupying bin *i*, *R_i_* is the mean firing rate for bin 𝑖, and *R* is the overall mean firing rate.

6. **Grid Score**

The hexagonal gridness measure is quantified using the Hexagonal Gridness Score (HGS) on the firing fields of each neuron. This measure identifies grid cells, which are neurons that exhibit a characteristic hexagonal firing pattern in spatial environments. The HGS is computed using the equations provided by [Hafting et al. (2005)].

**Autocorrelation Map Calculation**

The autocorrelation map 𝑟(𝜏𝑥, 𝜏𝑦) is calculated using the following formula:

$r\left( \tau x , \tau y \right)= \frac{M\sum_{x,y} \lambda\left( x,y \right)\lambda\left( x-\tau_{x},y-\tau_{y} \right)- \sum_{x,y} \lambda\left( x,y \right)\sum_{x,y} \lambda\left( x-\tau_{x},y-\tau_{y} \right)}{\sqrt{[M \sum_{x,y} \lambda\left( x,y \right)^{2}-{[\sum_{x,y} \lambda\left( x,y \right)]}^{2}]-[M\sum_{x,y} \lambda\left( x-\tau_{x},y-\tau_{y} \right)^{2}]-[\sum_{x,y} \lambda{(x-\tau_{x},y-\tau_{y})]}^{2}]}}$ In this equation:

- *r*(*τx*, *τy*) represents the autocorrelation map.
- *λ*(*x*, *y*) is the firing rate at the (𝑥,𝑦)(*x*,*y*) location of the rate map.
- *M* is the total number of pixels in the rate map.
- *τx* and *τy* correspond to the spatial lag in the *x* and *y* directions, respectively.

**Hexagonal Gridness Score Calculation**

The Hexagonal Gridness Score (HGS) is then determined using the following formula:

$$HGS=min\left[ cor\left( r,r^{{60}^{^{\circ}}} \right),cor\left( r,r^{{120}^{^{\circ}}} \right) \right] -\max\left[ cor\left( r,r^{{30}^{^{\circ}}} \right),cor\left( r,r^{{90}^{^{\circ}}} \right),cor\left( r,r^{{150}^{^{\circ}}} \right) \right]$$

In this equation:

- HGS stands for Hexagonal Gridness Score.
- 𝑟𝜃 is the autocorrelation map rotated by 𝜃 degrees.
- cor(⋅) stands for the correlation function.
- min(⋅) function returns the minimum of its two arguments, and max(⋅) returns the maximum of its arguments.
- The specific rotations used are 300°, 600°, 900°, 1200°, and 1500°.

These calculations provide a numerical measure of the hexagonal grid-like firing patterns of neurons, essential for identifying grid cells.

7. **Border Score**

To identify putative border fields, we first located clusters of neighboring pixels where the firing rates were above 0.1 times the maximum firing rate, with each cluster covering at least 200 cm². For experiments conducted in square or rectangular environments, we then estimated the field's coverage of a given wall as the fraction of pixels along the wall occupied by the field. The parameter $c_{M}$ was defined as the maximum coverage by any single field over any of the four walls of the environment.

We computed the mean firing distance *d_m_* by averaging the distances from all pixels within the field to the nearest wall, with the averaging process weighted by the firing rate. To do this, we normalized the firing rate by its sum over all pixels within the field, creating a distribution-like measure. We then normalized *d_m_* by dividing it by half of the shortest side of the environment, yielding a value between 0 and 1. The border score was defined by comparing *d_m_* to *c_M_*, the maximum coverage of any wall by a single field.

$$b= \frac{(c_{M}-d_{m})}{(c_{M}+d_{m})}$$

Border scores ranged from -1, indicating cells with central firing fields, to +1, indicating cells with fields that align perfectly with at least one entire wall. These scores provide insight into how fields expand across walls rather than away from them. The measure reaches saturation when the field's width approaches half the length of the environment. Cells with border scores above 0.5 were classified as 'border cells.' Only cells with stable border fields (spatial correlation > 0.5) were included in the analysis.

8. **Z-Score**

Responses to the object were quantified using z-scores. The z-score was calculated as

$$Z= \frac{(R_{obj}-R_{out})}{\left( \frac{sd}{\sqrt{N}} \right)}$$

Where, *R_obj_*, the mean firing rate within the area containing and immediately surrounding the object; *R*_out_, the mean firing rate outside the object location; *sd*, the standard deviation of the mean rate outside the object location; and *N*, the number of bins defining the object location. The area containing and immediately surrounding the object was defined as a 15 × 15 bin square (0.38 a.u * 0.38 a.u.) in the rate map, with the object's location at its center. We calculated the average firing rate from an equivalent number of bins randomly selected from the area outside the object location to determine the mean activity outside the object location.

For trace cell experiments involving object shifts, the R_obj_ was the mean firing rate of all the bins occupied by all the previous locations of the object and R_out_ stays the same. It gave a measure of increased cumulative firing rate for the older object positions compared to baseline firing outside object region.

9. **Path Integration (PI) layer**

This layer consists of oscillatory neurons that have one-to-one connection with neurons from the HD layer (Soman, Muralidharan, and Chakravarthy 2018). The process involved in this layer can be divided into two major stages. The first stage is frequency modulation of the HD responses. The second stage consists of low pass filtering to eliminate time-dependent high frequency oscillations, which gives the path integration output.

It is evident that theta oscillations exist in the Hippocampal formation and are modulated almost linearly with velocity (Burgess, Barry, and O’Keefe 2007; Geisler et al. 2007). The simplest way to represent frequency is in the form of sinusoids. Hence, we start with sinusoids with theta frequency ($\boldsymbol{\omega)}$ in eqn 1.

The first stage of frequency modulation (FM) of PI is described by the following equations:

${PI}_{i_{fm}}= sin[\omega t + \beta\int({HD}_{i})dt]$	(1)

$= sin[\omega t + \beta\int(v(t).u_{i})dt]$

$= sin[\omega t + \beta z.u_{i}]$	(2)

where, ω is the average base angular frequency of the oscillators which lies in theta frequency (4Hz – 8Hz) observed in the hippocampus, z is the displacement of the animal from its initial position, β is the scaling factor and $\boldsymbol{HD}_{\boldsymbol{i}}$ is the output of Head direction layer (methods section).

This was the model of path integration used in (Soman, Muralidharan, and Chakravarthy 2018; Soman, Muralidharan, and Chakravarthy 2018) (eqn. 1). This is one of the models which overcomes the constraint of specific direction tuning. However, it was also shown that with certain improvements, the OI models can also overcome this problem (Hasselmo and Shay 2014; Kropff and Treves 2008). Note that the PI cell described in eqn. (2) is a spatio-temporal model dependent explicitly on space, z, and on time, t. Such a model was successful in explaining certain temporal phenomena like phase precession (Soman, Muralidharan, and Chakravarthy 2018; Soman, Muralidharan, and Chakravarthy 2018). However, most spatial cells are described in purely spatial terms (e.g., place cells and grid cells), depicting their responses as exclusive functions of space. In the present study, we are interested in describing these purely spatial responses. Therefore, we convert the spatio-temporal model of eqn. (2) into a purely spatial model by an averaging process described below.

In order to eliminate temporal variation, a sin ($\boldsymbol{\omega t}$) the term is multiplied with the ${PI}_{i_{fm}}$ shown in eqn. 2, and passed through a low pass filter which blocks the high-frequency signals as shown in eqn. 3 and eqn. 4.

$PI_{i}=sin\left( \omega t \right) sin\left[ \omega t+\beta z.u_{i} \right]$	(3)

$PI_{i}=cos\left( \beta z.u_{i} \right)-cos(2\omega t+$ $\beta z.u_{i}$)

After eliminating the high-frequency term, we have,

$PI_{i}\approx cos\left( \beta z.u_{i} \right)$	(4)

10. **Random Shuffling Test**

*Place cells*: The random shuffling test for place cells involves reshuffling the firing rate of each neuron 400 times to generate a normal distribution of spatial scores and sparsity index. Both spatial scores and sparsity index of each neuron is compared with the 99^th^ percentile of their respective distribution to see if these metric scores are obtained by chance.

*Grid cells*: To test if the HGS score of a grid cell is more than expected by chance, we performed a random shuffling test by shuffling the firing points of each neuron randomly 500 times over the trajectory and calculated the 95^th^ percentile. Those neurons whose HGS is greater than the 95^th^ percentile were classified as the grid cells.

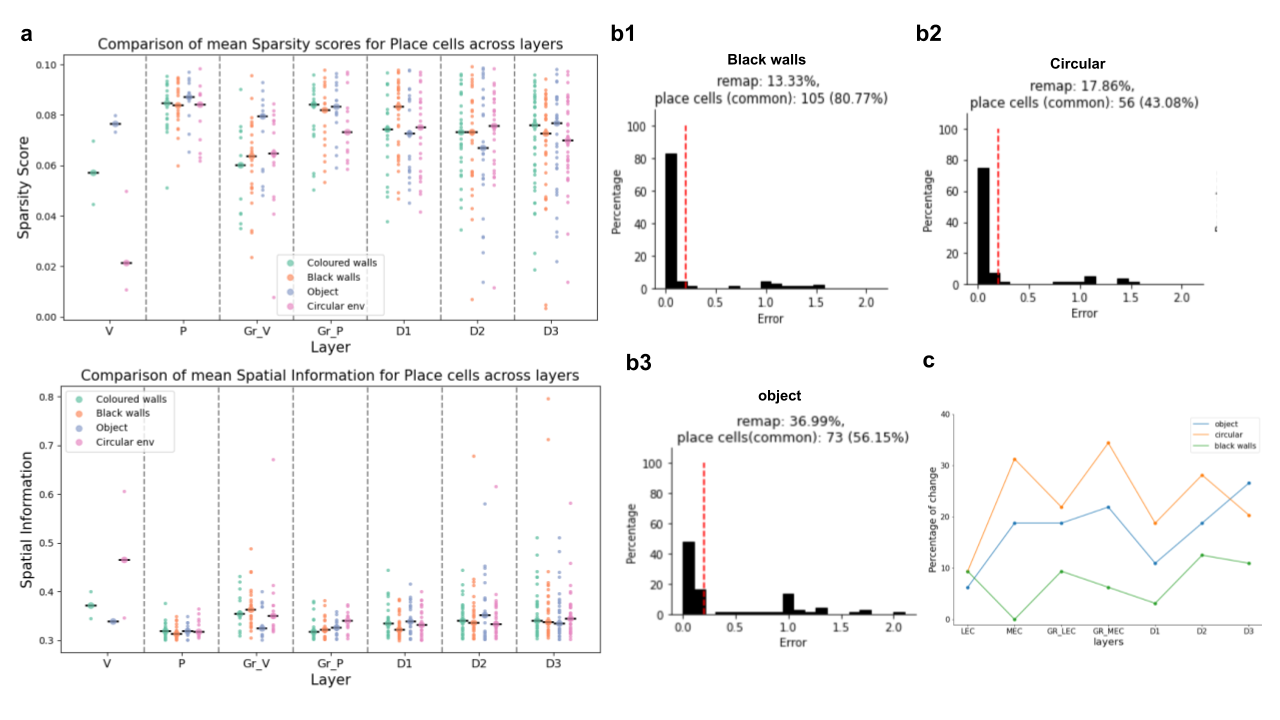

Figure S2: a) Comparison of mean sparsity scores (top) and spatial information (bottom) of place cells across all the layers in all the environments. b1, b2 and b3) shows the histogram of error (distance between the centroids place fields in the coloured walls environment and black walls, circular environment and environment with object respectively. The red dotted line represents the threshold of shift (10% of the size of environment with coloured walls) of the place field. The fields that shift more than the threshold are considered as remapped. C) percentage change of place cells across all the layers for each environment.

12. **Z-Score comparison with and without EWC:**

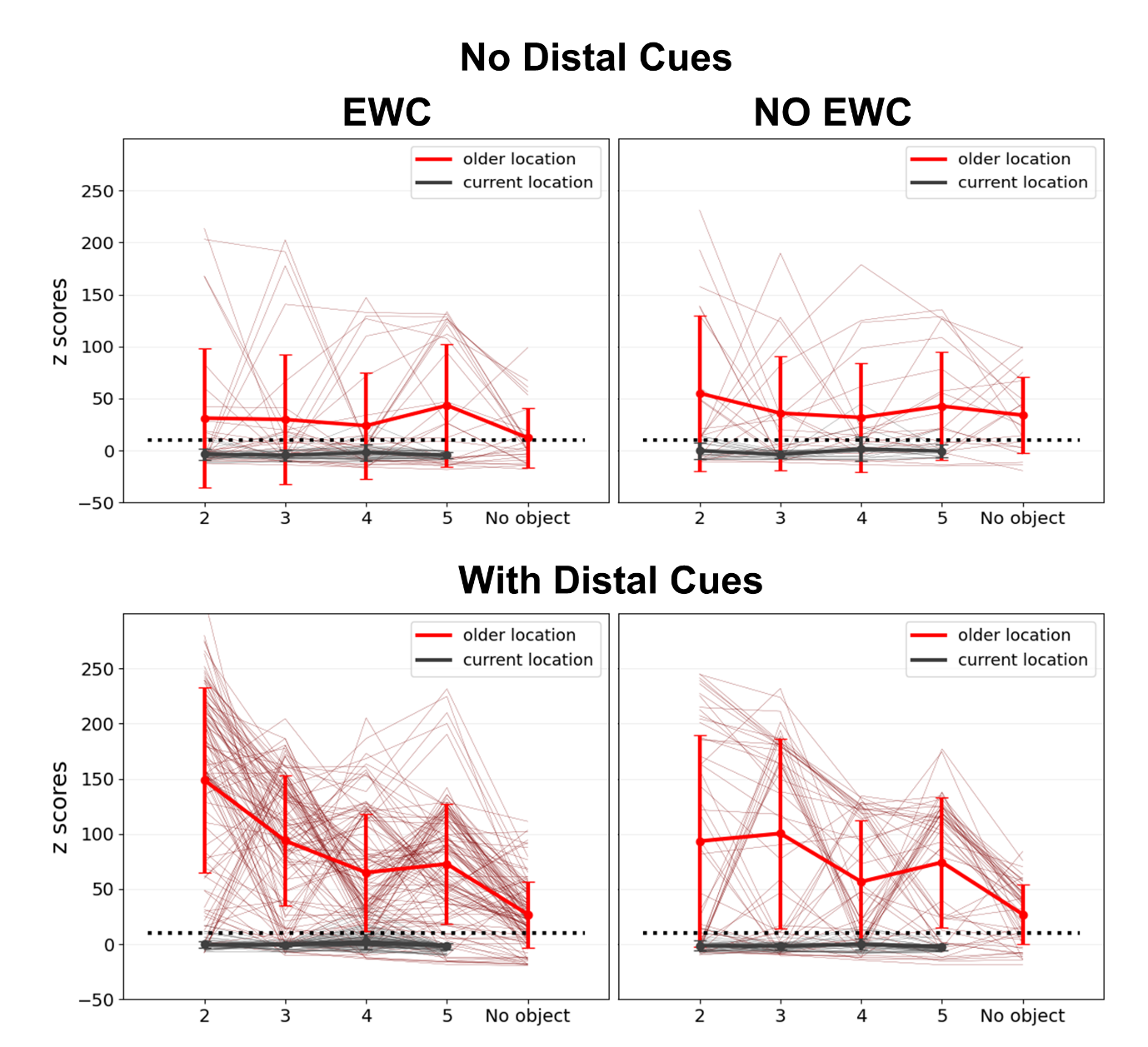

Figure S3: *Z scores of neurons with black walls (top row) and coloured walls (bottom row). The left figure in both the rows shows the z scores with Elastic Weight Consolidation (EWC) and the right figure shows z scores without EWC. Thick lines represent mean and thin lines represent z-score of individual neurons from session 2-6.*

13. **Trajectory Generation:**

Movement of the animal inside the square box is designed using the dynamics of

curvature-constrained motion. We used a dynamical system which is used to model

Dubins and Reeds-Shepp‘s cars (Takei and Tsai 2013). The dynamics are given as

follows:

$\dot{x}\left( t \right)= \sigma\left( t \right)cos\theta(t)$	(1)

$$\dot{y}\left( t \right)=\sigma\left( t \right)sin\theta(t)$$

$$\sigma\left( t \right)= r_{d}$$

$r_{d}= \left\| X_{pos}- X_{wall} \right\|$	(2)

$\left| \dot{\theta}(t) \right| \leq\frac{\gamma(t)}{\rho}$	(3)

where x and y are the 2D coordinates of the animal, $\theta$ Θ is the direction of movement of the animal at a given time, ρ>0 represents the minimum turning radius of the animal

and σ controls the speed of the animal. $X_{pos}$ and $X_{wall}$ are the Cartesian coordinates of

the position and the wall at which the animal is directing. $r_{d}$ is the distance from the animal’s current position to the nearest wall it encounters if it travelled continuously in that same direction. To ensure that the animal does not cross the boundary, the speed σ is set as $r_{d}$ and the animal’s speed considerably slows down as it nears the wall which is a reasonable assumption to consider for further simulations. Furthermore, in order to introduce the element of randomness in the animal’s trajectory a dynamic variable γ(t) is introduced in Eqn. (3). γ(t) ε [-1 1] is designed in such a way that there is high degree of randomness in the animal’s trajectory when the current position is far away from the walls compared to when it is close to it. A softmax rule is imposed for the probability of the animal to take a right or a left turn at a given instant. The parameter known as temperature in the softmax (here referred as α) is controlled by the distance of the animal to the walls. α is inversely proportional to the distance from the walls and its value becomes smaller and hence more random in picking the direction to move compared to when the distance to the walls is smaller and it sticks to a single direction. The dynamics of γ(t) is given by

$\dot{\gamma}\left( t \right)= -\gamma\left( t \right)+A$	(4)

$$Act \epsilon\left\{ 1, -1 \right\}$$

$$Act=sign[S_{max}\left( Q, \alpha\right)-rand$$

$$Q=[1 0.9]$$

$$\alpha= \frac{1}{r_{d}+c}$$

where Act is the action to be taken, Q is the value for each action, Smax is the Gibbs

softmax function which gives the probability of selecting an action which here is the

direction of motion and is given as:

$S_{max}\left( Q, \alpha\right)= p_{m}\frac{exp(\alpha Q)}{\sum_{i} exp(\alpha Q_{i})}$	(5)

$p_{m}$ is a parameter that switches discreetly between 1 and -1 after certain number of

iterations so that the animal can turn both directions closer to the walls.

**Analysis of grid cells that are losing their gridness across environments.**

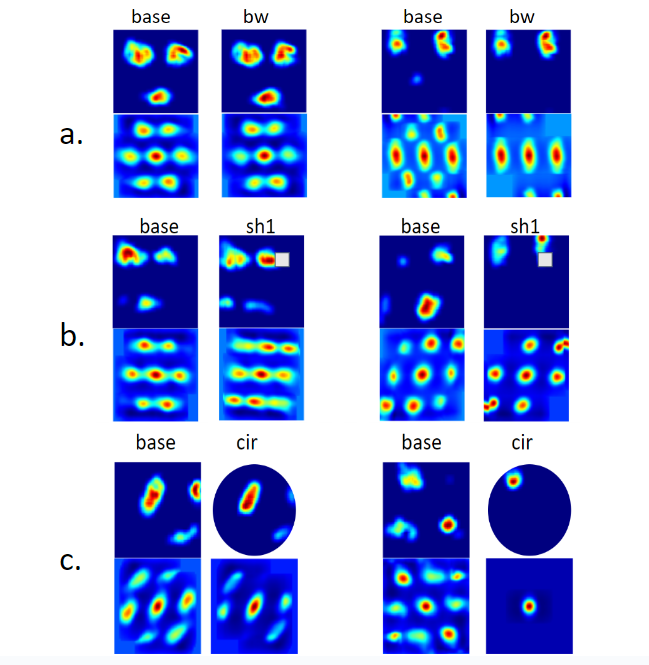

Figure S4: illustrates examples of hexagonal grid cells losing their gridness due to environmental modifications.

In each part of the figure (a, b, c), there are two sets of firing rate maps:

- The left set of each part shows hexagonal grid cells that lost their gridness because their auto-correlation rate maps shrank, despite their firing rate maps appearing similar.
- The right set in each part demonstrates hexagonal grid cells losing their gridness due to the disappearance of some place fields following environmental changes.

Specific environmental changes depicted are: a. Transition from an environment with colored walls without objects to one with black and white walls without objects. b. Change from an environment with colored walls without objects to colored walls with an object. c. Shift from an environment with colored walls without objects to a circular environment with colored walls and no objects.
